# Temporal weighting of cortical and subcortical spikes reveals stimulus dependent differences in their contributions to behavior

**DOI:** 10.1101/2023.08.23.554473

**Authors:** Jackson J. Cone, Autumn O. Mitchell, Rachel K. Parker, John H.R. Maunsell

## Abstract

The primary visual cortex (V1) and the superior colliculus (SC) both occupy stations early in the processing of visual information. They have long been thought to perform distinct functions, with V1 supporting perception of visual features and the SC regulating orienting to visual inputs. However, growing evidence suggests that the SC supports perception of many of the same visual features traditionally associated with V1. To distinguish V1 and SC contributions to visual processing, it is critical to determine whether both areas causally contribute to perception of specific visual stimuli. Here, mice reported changes in visual contrast or luminance near perceptual threshold while we presented white noise patterns of optogenetic stimulation to V1 or SC inhibitory neurons. We then performed a reverse correlation analysis on the optogenetic stimuli to estimate a neuronal-behavioral kernel (NBK), a moment-to-moment estimate of the impact of V1 or SC inhibition on stimulus detection. We show that the earliest moments of stimulus-evoked activity in SC are critical for detection of both luminance or contrast changes. Strikingly, there was a robust stimulus-aligned modulation in the V1 contrast-detection NBK, but no sign of a comparable modulation for luminance detection. The data suggest that perception of visual contrast depends on both V1 and SC spiking, whereas mice preferentially use SC activity to detect changes in luminance. Electrophysiological recordings showed that neurons in both SC and V1 responded strongly to both visual stimulus types, while the reverse correlation analysis reveals when these neuronal signals actually contribute to visually-guided behaviors.

## Introduction

In the mammalian visual system, information leaving the eye is routed to two distinct processing centers, the primary visual cortex (V1), via projections from the lateral geniculate nucleus (LGN), and the superior colliculus (SC) in the rostral midbrain. V1 serves as the entry point to the cortical visual hierarchy where neuronal activity corresponds to fundamental visual features (e.g., edges, orientation, motion direction) that form the foundations of more complex percepts.^1–3^ As V1 represents the final bottleneck before visual information is distributed to multiple higher-order cortical visual areas^4^, disruption of V1 function dramatically impairs visual perception across species.^5–9^ In contrast, the SC has long been studied for its highly conserved role in orienting movements of the body, head and eyes.^10–15^ More recently, the SC has also been shown to contribute to covert orienting behaviors like visual spatial attention.^16^ For example, inactivation of the SC produces deficits in selecting which of several competing stimuli will inform perceptual judgements, even in the absence of overt orienting responses.^17^ Taken together, these observations suggest a division of labor where V1 contributes to perception and recognition of visual stimuli while the SC is involved in the selection of visual targets.^18^

Accumulating evidence increasingly challenges these divisions. Work in both rodents and primates suggests that, like V1, activity of SC neurons is used to inform perception of visual features. Mouse SC neurons are tuned to many of the same visual inputs as their V1 counterparts, including motion direction,^19,20^ orientation,^21,22^ luminance^23^, and contrast.^24^ Furthermore, SC responses are largely preserved following removal of V1, arguing that visual selectivity in the SC is not simply inherited from cortical input.^19^ Behavioral evidence further suggests that SC visual representations contribute to perception. Optogenetic inhibition of mouse SC impairs detection of orientation changes most strongly when delivered during a 100 ms window immediately following the change, suggesting that SC activity primarily contributes to sensory evaluation in this task.^25^ Furthermore, chemogenetic inactivation of one genetically defined subtype of SC neuron impairs prey detection, while inactivation of a different subtype disrupts orienting towards said prey.^26^ While a much larger proportion of retinal ganglion cells sends input to the SC in mice (∼85%)^27^ compared to primates (∼10%),^28^ data from primates similarly support a role for SC in sensory evaluation. Recent data suggests that magnification of the foveal representation in monkey SC is comparable to that in V1.^29^ Moreover, in addition to deficits in target selection in the presence of distractors, SC inactivation also impairs judgements of visual motion when distractors are omitted.^17,30^ Thus, the evaluation and perception of visual stimuli may use both the SC and V1 in a cooperative manner.

The extent of cortical and subcortical contributions to visually guided behaviors might vary depending on the nature of visual input. For example, V1 activity may be weighted more strongly for perception when visual stimuli are better matched to V1 compared to SC receptive fields (i.e., contrast versus luminance). Addressing these questions requires methods capable of distinguishing neuronal contributions to perception, sensorimotor coordination, and/or action execution for different visual stimuli. We recently adapted reverse correlation methods for optogenetic inhibition of visual processing in behaving mice.^31^ Mice performed a visual detection task while headfixed. On a subset of trials, randomized patterns of optogenetic stimulation were delivered to inhibitory interneurons in V1. Reverse correlating the optogenetic stimuli with task outcomes (e.g., hit, miss) yielded a neuronal-behavioral kernel (NBK), a moment-to-moment estimate of the periods of V1 activity that were used to guide behavior. Here, we leveraged optogenetic reverse correlation in the SC or V1 of mice as they reported changes in visual contrast or luminance to explore which periods of activity – and in which brain areas – were used for guiding behavior. We show that the earliest moments of spiking activity in SC and V1 are differentially weighted for guiding behavior in a stimulus-dependent fashion, suggesting that both cortical and subcortical signals coordinate perception of some elementary visual features but not others.

## Materials and Methods

### Animal Preparation

All animal procedures followed NIH guidelines and were approved by the Institutional Animal Care and Use Committee of the University of Chicago. We used mice that were heterozygous for Cre recombinase in Parvalbumin (PV) expressing cells (V1 behavioral experiments, 8 mice, 4 female) or glutamic acid decarboxylase 2 (GAD2) expressing cells (SC behavioral experiments, 11 mice, 6 female). These strains provide a targeting specificity of >92% for PV+ or GAD2+ neurons.^32–34^ Animals were outbred by crossing homozygous Cre-expressing mice (PV: JAX Stock #017320; SC: JAX Stock #010802) with wild-type BALB/c mice (JAX stock #000651). Animals were singly housed on a reverse light/dark cycle with *ad libitum* access to food. Mice were water scheduled throughout behavioral experiments, except for periods around surgeries.

Mice (2–4 months old) used for SC experiments were implanted with a titanium headpost while mice used for V1 experiments received a headpost and a cranial window to give stable optical access to cerebral cortex.^35,36^ For surgery, animals were anesthetized with isoflurane (induction, 4%; maintenance 1.0-1.5%) and given ketamine (40 mg/kg, IP) mixed with xylazine (2 mg/kg, IP). Body temperature was maintained with a heating pad. The headpost was secured to the skull with acrylic (C&B Metabond, Parkell) using aseptic technique. To implant the optical window, a craniotomy was made over V1 in the left cerebral hemisphere (3.0 mm lateral and 0.5 mm anterior to lambda) and covered with a glass window (3.0 mm diameter, 0.8 mm thick; Tower Optical). Mice were given analgesics postoperatively (buprenorphine, 0.1 mg/kg and meloxicam, 2 mg/kg, IP).

To localize V1 underneath the optical window, we measured changes in intrinsic autofluorescence using visual stimuli and epifluorescence imaging.^37^ Autofluorescence produced by blue excitation (470 ± 40 nm, Chroma) was captured using a long-pass filter (500 nm cutoff), a 1.0X air objective (Carl Zeiss; StereoDiscovery V8 microscope; ∼0.11 NA) and a CCD camera (AxioCam MRm, Carl Zeiss; 460 x 344 pixels; 4 x 3 mm FOV). The visual stimuli were full contrast drifting sinusoidally modulated luminance gratings appearing through a two-dimensional Gaussian aperture (Gabors). Gabor stimuli (10° SD; 30°/s; 0.1 cycles/deg) were presented for 10 s followed by 6 s of mean luminance at 1 of 5 visual field locations. The response to each visual stimulus was computed as the fractional change in fluorescence during the first 8 s of the stimulus presentation relative to the last 4 s of the preceding blank.

### Behavioral Task

After recovery from headpost and/or window implantation surgery, mice were trained to manipulate a lever to respond to changes in a visual display for a water reward.^38^ During behavioral sessions, mice lay in a sled with their head fixed. Mice were positioned in front of a calibrated visual display that presented a uniform gray field. Mice self-initiated trials by depressing the lever, and a neutral tone indicated the start of each trial. The prestimulus period was randomly drawn from a uniform distribution (500-3000 ms), after which a static achromatic visual stimulus appeared (*contrast*: odd-symmetric Gabor, σ = 5-7°,0.1 cycles/deg; *luminance*: gaussian filtered black spot, σ = 5-7). Mice had to release the lever within 100-700 ms after stimulus onset to receive a liquid reward (1-4 µL). Trials in which mice failed to release the lever within the response window resulted in a brief time out (1500-2000 ms). Trials in which mice released the lever before the onset of visual stimuli (i.e., false alarms) were unrewarded and excluded from analyses related to visual processing (Figure 2). False alarm trials were analyzed separately to assess the impact of optogenetic stimulation of motor responses (Figure 3). Task parameters were slowly adjusted over several weeks until mice responded reliably to visual stimuli (σ = 5-7°) in the right hemifield near the location of the V1/SC retinotopic map where optogenetic stimulation would be delivered during testing sessions.

### AAV injections

After achieving expert performance (i.e., d’ > 2.0 over multiple sessions, median reaction times < 350 ms) mice were injected with Cre-dependent AAV viruses carrying ChR2-tdTomato.^39^ Mice were anesthetized (isoflurane, 1.0-1.5%) and their head post was secured in a mount. We used a volume injection system (World Precision Instruments UMC4) to deliver two 200 nl boluses of AAV9-Flex-ChR2-tdTomato (∼10^12^ viral particles; Penn Vector Core) at a rate of 50 nl/min. The injector was left in position at the injection site for 5 minutes before and after each injection.

Injections were targeted to retinotopically defined regions of V1 or SC using different approaches. For V1 injections, we targeted the monocular region of V1 based on each animal’s retinotopic map (25° azimuth; ±15° elevation) and injected a region of V1 that corresponded to the selected location for visual stimuli presented on the visual display during subsequent behavioral sessions. The cranial window was removed using aseptic technique. The first injection site was 400 µm below the cortical surface. After the first injection, a second injection was made 200 µm below the cortical surface.. The injector was retracted and a new cranial window was then sealed in place. Once Td-Tomato fluorescence was visible underneath the optical window (2-3 weeks post-injection) we attached an optical fiber (400 µm diameter; 0.50 NA; ThorLabs) above ChR2-expressing neurons within 500 µm of the cranial window (∼1.3 mm above the cortex).

For SC virus injections, we performed electrophysiological recordings to map retinotopy at the injection site (*see Electrophysiological Recordings and Analysis*). After induction with isoflurane, the metabond over bregma was carefully drilled away until bregma could be clearly visualized. We then drilled a small burr hole over the left SC (3.5 mm posterior and 1.8 mm lateral to bregma) and lowered a 4-channel electrode (CQ4 Probe, Neuronexus) into the superficial SC (−1.8 mm below the surface of the brain). After identifying multiunit activity on the electrode, the electrode was allowed to settle for 30 minutes. A gamma corrected visual display was then positioned in the visual hemifield opposite of the recording site (∼10 cm viewing distance) and the eyes were kept moist with 0.9% saline throughout the session. Electrode signals were amplified, bandpass filtered (750 Hz to 7.5 kHz) and sampled around threshold voltage crossings (Blackrock, Inc.). Counterphasing luminance patches (2D Gaussian, ∼15° σ) were presented at various locations on the visual display while multi-unit spiking responses were monitored on an audio amplifier. After identifying the spatial receptive field location at the SC recording site, the electrode was retracted, and an injector was lowered into the brain and advanced an additional 400 µm beyond the recording location (∼ −2.2 mm DV from brain surface). Following the first injection, the injector was raised 400 µm and a second injection began. The injector was then retracted and an optical fiber (200 µm diameter; 0.50 NA; ThorLabs) was lowered to 50 µm above the second injection site. The burr hole was filled with Kwik-Cast silicone sealant (World Precision Instruments) and the optical fiber was fixed in place with metabond. Mice were then removed from the headpost holders and returned to their home cage for recovery.

### Optogenetic Stimulation and Alignment with Visual Stimuli

Behavioral sessions with optogenetic stimulation began >3 weeks after ChR2 injection. Light was delivered through the optic fiber using a power-calibrated LED source (455 nm; ThorLabs). To prevent mice from seeing optogenetic stimuli, we enclosed the fiber implant in blackout fabric (ThorLabs) attached to the head post using a custom mount. The timing of the optogenetic stimulus delivered on each trial was synchronized with that of the visual stimulus using a photodiode mounted on the video display.

In preliminary experiments, we optimized the alignment between the retinotopic location of the visual stimulus presented on the visual display with the site of optogenetic perturbations in SC or V1. During these sessions (∼5-7 sessions/mouse), the visual stimulus (Gabor, σ = 5-7°) alternated randomly between one of two locations in the right hemifield. The location of visual stimuli were offset by 5-10°, either in azimuth or elevation within an individual session. On a random half of trials, visual stimuli were paired with optogenetic stimulation of inhibitory neurons in SC or V1 (single square pulse, 0.5 mW, onset/offset concurrent with the visual stimulus). For each visual stimulus location, we computed the change in the hit rate for trials with versus without optogenetic stimulation to determine the visual stimulus location that was most affected by optogenetic perturbation. The visual stimulus location that produced the largest decrease in detection performance on trials with optogenetic stimulation compared to trials without was used for all subsequent experiments. Data from these sessions is summarized in Figure 1B but were otherwise not included in our primary analyses. For all remaining behavioral sessions, the visual stimulus was presented at the optimal location for each mouse identified in these preliminary experiments (except where specified).

**Figure 1.**
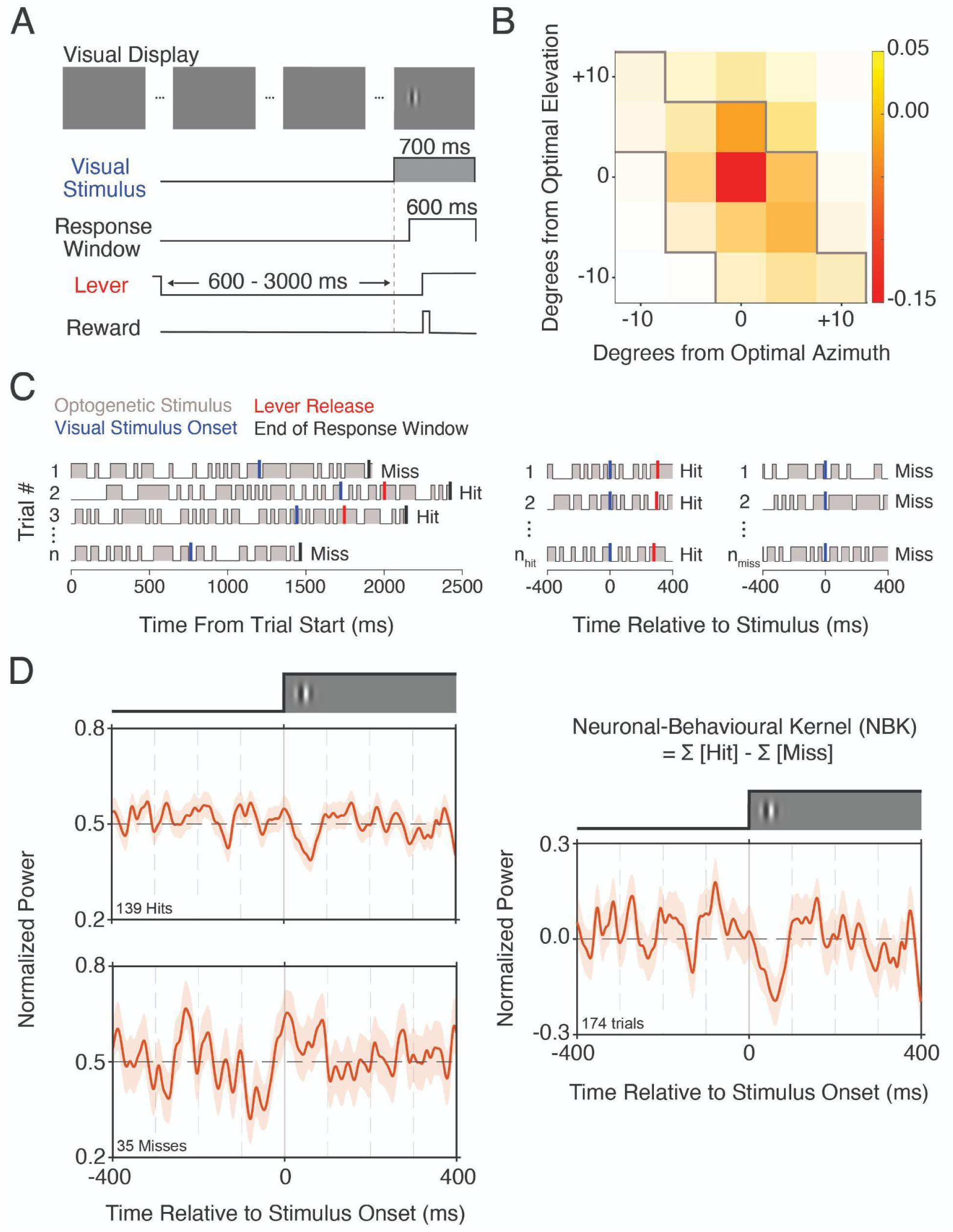
Task design, optimization, and construction of the neuronal-behavioral kernel. **A)** Trial schematic of the visual detection task. The mouse was required to release a lever shortly after a small visual stimulus (contrast or luminance patch) appeared on the display. The visual cstimulus appeared at a random time in each trial. **B)** Heatmap quantifies the effect on detection performance of optogenetic stimulation (0.5 mW, square pulse) delivered concurrently with the visual stimulus as a function of visual stimulus azimuth (x-axis) and elevation (y-axis). Hue depicts impairment or enhancement in detection performance while saturation depicts the relative number of trials collected at each visual stimulus location. Sessions in which stimulation resulted in a reduction in hit rate lie within the gray bounds. **C)** (left) Representative optogenetic stimulus profiles taken from individual trials in a session. For each trial, gray indicates the 25 ms periods randomly assigned for optogenetic stimulation, blue lines mark visual stimulus onset, red lines mark lever releases, and black lines mark the end of the reaction time window. Outcomes are indicated to the right of each trial. (middle) An optogenetic kernel based on hit trials was constructed aligning the optogenetic stimulus profiles from the 139 trials with successful detections to the onset of the visual stimulus. (right) An optogenetic kernel was similarly constructed from the 35 trials on which the mouse failed to respond to the stimulus. **D)** (top left) The optogenetic kernel constructed from the 139 hit trials. A brief dip in the kernel is evident 50 ms after visual stimulus onset (t = 0) shows that successful detections were more likely when the optogenetic stimulation was weaker during this period. (bottom left) The optogenetic kernel constructed from the 35 miss trials. The brief rise in the kernel immediately after the visual stimulus onset shows that misses were more likely when the optogenetic stimulation was stronger during this period. (right) A full NBK was constructed using hit and miss trials by subtracting the mean power and inverting the optogenetic stimulus profiles from the miss trials before averaging. The shaded regions indicate ±1 SEM. Cortical and subcortical visual areas make distinct contributions to perception of simple visual stimuli.

### Behavioral Analysis

The proportion of successful lever releases is a useful measure of animal performance, but fails to account for guessing. The d’ measure from Signal Detection Theory^40^ addresses this issue and provides a more complete measure of the animal’s discrimination abilities. d’ is computed using both hit and false alarm rates, but the false alarm rate is not directly available in the visual detection task. We estimated the false alarm rate for each session by dividing the total trial time available for early lever releases by the number of early releases. The trial time available for early releases included the trial time before each early release and the time before each stimulus presentation on other trials (accounting for the ∼100 ms period after stimulus onset during which responses were considered early because they were too fast to be valid). The rate of early lever releases was multiplied by the duration of the response window to obtain the overall probability of a false alarm. The false alarm probability was then used to convert the raw hit rate to a true hit rate by removing the fraction of hits attributable to spontaneous lever releases during the response window (false hits). A d’ value was computed separately for trials with and without optogenetic stimulation for each session. The rate of false alarms was indistinguishable betweens trials with versus without optogenetic stimulation (see *Behavioral performance*) so we used the combined false alarm rate when computing d’ for stimulated and unstimulated trials.

### White Noise Optogenetic Stimulation and Analysis

During experiments using white noise optogenetic stimulation, optogenetic stimulation was delivered on a random half of all trials with the 30% contrast stimulus. We also included “top-up” stimuli (50% contrast) that were never paired with optogenetic stimulation in order to estimate a lapse rate for each session. On trials with optogenetic stimulation, the optogenetic stimulus was delivered throughout the trial. The power of the optogenetic stimulus ramped up to the average mean power over the first 250 ms of the trial to avoid abrupt changes in neuronal activity. The ramp period was excluded from analysis. The mean power used for optogenetic stimulation was capped at 0.25 mW, but needed to be varied across mice to keep the behavioral disruption modest (median: 0.15 mW, range 0.09 - 0.25 mW), likely owing to differences in the strength and spatial distribution of virus expression, optic fiber alignment, and behavioral capacity. Binary white noise optogenetic stimuli were generated by randomly assigning zero or twice mean power to successive 25 ms bins with equal probability (phase randomized on each trial). The resulting optogenetic stimulus had equal power across all frequencies represented and is, therefore, a quasi-white noise stimulus.^41^ Optogenetic stimulation at substantially higher frequencies would have been filtered by the kinetics of the ChR2^42^ (∼15 ms decay time) and the PV-principal neuron synapse^43^ (∼15 ms rise and decay times), and would have reduced the power in the relevant portion of the spectrum.

A first-order Wiener kernel was calculated from the optogenetic stimuli for trials that ended with a hit or a miss. The optogenetic stimulus profiles trials were normalized, aligned to the onset of the visual stimulus, profiles from miss trials were inverted, and then averaged for all trials (V1 contrast: 21,789 hit trials, 10,851 miss trials, median = 27 sessions, range 20-37 sessions; SC contrast: 20,412 hit trials, 11,169 miss trials, median 27 sessions, range 10-52 sessions; V1 luminance: 22,146 hit trials, 8,839 miss trials, median 28 sessions, range 12-31 sessions; SC luminance: 34,767 hit trials, 18,205 miss trials, median 29 sessions, range 24-55 sessions). Kernels were low-pass filtered with a corner frequency of 90 Hz to eliminate noise beyond the frequency range of the optogenetic stimulus. Data from different animals were combined by normalizing the maximum optogenetic power to 1. Confidence intervals were generated using a bootstrap procedure with 10,000 draws with replacement.

Our prior work shows that the magnitude of the NBK scales with the effect of the optogenetic stimulation on performance measured as the change in d’ for trials with versus without optogenetic stimulation (Day-Cooney, Cone, and Maunsell 2022). Thus, we only considered sessions where Δd’ was ≤ +1 SD from the mean across mice to omit sessions in which optogenetic stimulation did not interfere with detection performance. The Δd’ distributions were negatively skewed, so this criteria excluded only a small number of sessions with positive Δd’ (V1 contrast: 4% (9/218 sessions); SC contrast: 4% (8/193 sessions); V1 luminance: 7% (12/163 sessions); SC luminance: 6% (23/390 sessions)).

In a separate analysis, we computed an NBK after aligning the optogenetic stimuli to false alarms to determine whether those errors are associated with optogenetically driven fluctuations in spiking activity in V1 or SC (Figure 3A-B). Finally, we computed an NBK using optogenetic stimuli aligned to the onset of behavioral responses (Figure 3C-D), to explore neuronal contributions that were temporally aligned with correct lever releases following the onset of visual stimuli. These NBKs were computed using these same sessions that were included above.

Six of eight V1 mice (2 female) and seven of 11 SC mice (2 female) performed both contrast and luminance detection sessions. Five of eight V1 mice, including four that performed both contrast and luminance detection sessions, and six of 11 SC mice, including four that performed contrast and luminance detection sessions, also conducted sessions where the visual stimulus was deliberately moved out of retinotopic alignment with the optogenetic stimulus (see Figure S3). There were no exclusion criteria applied to these sessions but there is no expectation that optogenetic stimulation offset from the visual stimulus representation should impair detection performance.

### Electrophysiological Recordings and Analysis

We made acute electrophysiological recordings from single and multiunit sites in V1 or SC in awake, head fixed mice while they passively viewed visual stimuli. Before recording sessions, naive mice were surgically prepared as described above (i.e., headpost, window). All mice were injected with AAVs at least 3 weeks before recordings to ensure expression of ChR2 in PV+ or GAD2+ neurons in V1 or SC, respectively. We used linear single-shank, multi-contact electrode arrays (Poly2 configuration, NeuroNexus Inc.) that were custom made with a 200 μm core optical fiber (0.22 NA) that terminated 50 μm above the most superficial electrode contact. Electrodes were electroplated with a gold solution mixed with carbon nanotubes to impedances of 200-500 kΩ before all recording sessions.^44,45^

Mice (V1: n = 8 mice, 4 female; SC: n = 11 mice, 4 female) were anesthetized with isoflurane (1.2–2.0% in 100% O_2_) and head fixed. A gamma corrected visual display was positioned in the visual hemifield opposite of the recording site (∼10 cm viewing distance) and the eyes periodically moistened with 0.9% saline throughout recording sessions. For V1 recordings, we visualized ChR2-expressing areas of V1 by imaging tdTomato fluorescence with a fluorescence microscope and camera (Zeiss). The cranial window was then removed and linear multielectrode arrays lowered into V1 through a slit in the dura. The craniotomy was then covered with 3% agar (MilliporeSigma) dissolved in aCSF (TOCRIS). The agar was covered with silicone oil (60,000 centistokes, MilliporeSigma) to prevent drying. For SC recordings, the burr hole above SC marking the original injection site was enlarged and the electrode was lowered such that the contacts spanned superficial and intermediate SC (2.4 to 1.6 mm below skull surface). After a 1 hour recovery period, anesthetic was removed and there was an additional 1 hour wait before the start of data collection.

We first sought to characterize the average STAs from populations of SC or V1 neurons in response to white noise optogenetic stimuli (0.25 mW mean power). For this dataset, we sampled from 16 unique electrode positions in V1 of 8 mice (4 female) and 21 unique locations in the SC of 11 mice (6 female). During each trial, white noise optogenetic stimuli were delivered to V1 or SC together with a full-screen counterphasing gabor stimulus (temporal frequency 2 Hz) presented on the visual display. The visual stimulus was present throughout every trial and simply served to elevate spiking activity above baseline rates. In other recording sessions, we presented the same visual stimuli used during behavioral sessions (e.g., contrast or luminance) where a random half of all presentations also included white noise optogenetic stimulation of ChR2 expressing GABAergic neurons in V1 or SC.

In a different set of recordings, we assessed whether optogenetic stimulation of one structure (i.e., V1) impacted neuronal responses in the other brain area (SC). For SC recording sessions with V1 optogenetic stimulation (n = 2 mice, 1 female), we used PV-Cre mice that had been previously injected with ChR2 in V1. A burr hole was made over SC, the electrode was lowered into place, and the retinotopy of the recording site was manually confirmed as described above. After identifying the retinotopic location of the recording site, an optical fiber was secured in an adjustable mount and positioned over the corresponding ChR2-expressing retinotopic location of V1 using Td-Tomato fluorescence and V1 retinotopic mapping data. For V1 recording sessions with SC optogenetic stimulation (n = 3 mice, all males), we used Gad2-Cre mice that had previously been injected with ChR2 in the SC and an optical fiber had been affixed above the injection site. A small craniotomy was made over V1 and the electrode was positioned in the same retinotopic location in V1 as the site of SC ChR2 injections. We also performed recordings in V1 of mice that did not express ChR2 (n = 2 mice, 1 female). All mice had previously been implanted with a headpost and an optical window prior to recordings. The electrode was lowered into V1 which was identified underneath the window before the recording session with intrinsic signal imaging.

Delivery of stimuli and data acquisition were computer controlled. Concurrent visual and optogenetic stimuli were similar to those used during behavioral experiments (contrast stimuli: a Gabor patch with SD 10°; luminance stimuli: black spot spanning 30° with 500 ms duration). We recorded >200 repetitions of each stimulus condition (visual stimulus, visual stimulus + white noise optogenetic stimulation). Electrode signals were amplified, bandpass filtered (750 Hz to 7.5 kHz) sampled around threshold voltage crossings (Blackrock, Inc.) and spikes were sorted offline (OfflineSorter, Plexon, Inc.). Analyses were done using MATLAB (MathWorks) and custom code. We recorded both single and multi-units but did not differentiate between them because our primary interest was how optogenetic manipulations affected V1 and SC neuronal populations.

Units were considered visually responsive to visual stimuli (without optogenetic stimulation) if they exhibited a significant change in their average firing rate (p < 0.05; Wilcoxon signed-rank test) during the 10-110 ms after stimulus onset relative to the average firing rate during the 10-110 ms before stimulus onset. Units were defined as optogenetically responsive if there was a significant difference (p < 0.05; Wilcoxon signed-rank test) in the trial averaged firing rate between trials with and without optogenetic stimulation.

We calculated the spike triggered average (STA) optogenetic stimulus based on standard approaches.^46^ The power used for binary optogenetic stimuli was first normalized from −1 to 1. For each unit, optogenetic stimuli in windows 350 before to 100 ms after individual spikes occurred were extracted and averaged. Only units for which the minimum or maximum of the spike triggered average exceeded 8 SEM were used in further analyses. Eliminating the inclusion criteria did not qualitatively impact our observations.

Data are accessible upon reasonable request.

## Results

### Behavioral performance

Mice performed a visual detection task (Figure 1A) in which they reported the onset of a small contrast or luminance stimulus that appeared on a visual display after a random delay from trial start (Figure 1A).^8,38^ The task featured three trial types. The minority of trials (25-40%) were 50% contrast stimuli that were always presented without optogenetic stimulation. These trials were used to estimate lapse rates and offset reward losses resulting from optogenetic perturbations. The remaining trials were 30% contrast stimuli presented alone or together with white noise optogenetic stimulation. Trials without optogenetic stimulation enabled us to measure unperturbed performance at the tested stimulus intensity and estimate the effect of optogenetic stimulation on detection performance. We used a single visual stimulus intensity for estimating optogenetic-behavioral relationships as our prior work showed that optogenetic-behavioral kernels are visual stimulus dependent.^31^ We used signal detection theory to compute behavioral sensitivity (d’) and criterion (c) for the different visual stimulus conditions (see *Behavioral Analysis*). Summary statistics for d’ and c appear in *Supplementary Information*. Overall, mice performed the task well with low to moderate lapse and false alarm rates, resulting in high d’s and moderately conservative criterions (30% contrast without optogenetic stimulation; median d’ and c: V1 contrast: 2.53, 0.74; SC contrast: 2.16, 0.65; V1 luminance: 2.70, 0.72; SC luminance: 2.23, 0.64).

Figure 1B (as well as prior work^8,25,47^) shows that activation of GABAergic neurons in V1 or SC impairs performance in perceptual tasks. We computed the within-session change in d’ (Δd’) for 30% contrast stimuli presented with versus without white noise optogenetic stimulation. As expected, optogenetic stimulation of V1 or SC GABAergic neurons produced a small but significant reduction in detection performance for both visual stimuli (median Δd’ across all sessions: V1 contrast: −0.05; SC contrast: −0.05; V1 luminance = −0.06; SC luminance: −0.08, difference from 0, all p < 0.001, Wilcoxon signed-rank test). Trials with optogenetic stimulation also produced a small but significant elevation in criterion (median Δc across all sessions: V1 contrast: 0.03; SC contrast: 0.02; V1 luminance = 0.03; SC luminance: 0.04, difference from 0, all p < 0.001, Wilcoxon signed-rank test). The small changes in performance we observed are consistent with the random nature of the optogenetic stimulus as there are likely many trials in which the optogenetic stimulus would not be well aligned with the moments of neuronal activity that the mice were using to perform the task. Furthermore, the power delivered on trials with optogenetic stimulation was kept low to study these circuits as close to their normal operating ranges as possible (mean power on stimulation trials: V1 contrast = 0.11 mW, IQR: 0.07 - 0.14; SC contrast = 0.11 mW, IQR: 0.07 - 0.18; V1 luminance = 0.19 mW, IQR: 0.12 - 0.26; SC luminance = 0.08 mW, IQR: 0.06 - 0.10).

Notably, optogenetic stimulation had no detectable effect on the rate of false alarms (median probability of false alarm for 30% contrast visual stimuli, V1 contrast: visual = 0.02, IQR 0.01-0.03; visual + opto = 0.02, IQR 0.01-0.03; V1 luminance: visual = 0.02, IQR 0.01-0.03; visual + opto = 0.02, IQR 0.01-0.03; SC contrast: visual = 0.04, IQR 0.03-0.06; visual + opto = 0.05, IQR 0.03-0.06; SC luminance: visual = 0.04, IQR 0.02-0.07; visual + opto = 0.04, IQR 0.02-0.07; all p > 0.05, Kolmogorov-Smirnov test). A reduced hit rate with no change in false alarm rate is expected if the primary effect of stimulating inhibitory neurons in V1 or SC on random trials is to reduce the strength of the sensory signal. Lastly, our prior work shows that the magnitude of the NBK in V1 scales with Δd’.^31^

### White noise optogenetic inhibition

Delivering weak, stochastic optogenetic stimulation of inhibitory neurons (e.g. binary white noise optogenetic stimulation) enables efficient and unbiased sampling for how different moments of neuronal activity contribute to behavior.^31^ Following the optimization sessions discussed above, we switched to sessions in which white noise optogenetic stimulation was presented to ChR2-expressing V1 PV or SC GAD2 GABAergic neurons throughout a randomly selected subset of trials. We constructed the optogenetic stimulus train for each trial by randomly assigning zero or full power to each 25 ms trial interval (binary white noise). We selected 25 ms optogenetic stimulus intervals because the onset and offset kinetics of ChR2 activation sets an upper bound on the temporal resolution of this approach.^42^ Behavioral outcomes on trials with white noise optogenetic stimulation were used to construct a NBK (*see Materials and Methods*).

Figure 1C-D illustrates the analysis for an individual V1 stimulation session that produced a clear kernel. We first extracted all stimulated trials that ended in either a successful detection (hit) or a failure to detect (miss; Figure 1C). For the primary analysis, we separated hit and miss trials, aligned the trials on the time of visual stimulus onset, and normalized the binary optogenetic stimulus waveform for each trace to values of zero and one to account for different powers used across mice with different levels of virus expression (average maximum optogenetic power across sessions 0.25 mW, IQR = 0.20–0.30 mW; Fig. 1C middle, right). The visual stimulus-aligned optogenetic waveforms were then averaged to produce kernels separately for the hit and miss trials. Figure 1D shows the hit (top) and miss (bottom) kernels for this session. The average of the optogenetic traces remains near the mean normalized power (0.5), but there is a visible dip ∼50 ms after stimulus onset on hit trials (Figure 1D, top). This shows that reduced optogenetic stimulation of V1 PV+ neurons during this period was associated with a greater probability of visual stimulus detection. We expect a negative peak because lower optogenetic power decreases inhibition onto V1 excitatory neurons. Correspondingly, Figure 1D (bottom) shows that the miss kernel has greater power during the same interval indicating that greater optogenetic stimulation powers during this period were associated with a greater likelihood of failing to detect the stimulus. We combined the hit and miss trials from each session into a single NBK by averaging the mean-subtracted normalized optogenetic waveforms from all trials after inverting those from miss trials (first-order Wiener kernel; Figure 1D, right). The NBK represents the linear temporal weighting for how optogenetically activating V1 PV+ neurons at times relative to the visual stimulus onset relates to behavioral detection of that stimulus.^48^ Here, a normalized power of zero corresponds to no systematic effect of the optogenetic stimulus on detection performance. The large negative peak in the right panel of Figure 1D ∼50 ms after visual stimulus onset corresponds to reduced optogenetic stimulation at this time making the animal more likely to detect the visual stimulus. Smaller peaks at other times were not consistent across sessions.

Our primary goal was to use white noise optogenetic stimulation to explore moment-to-moment relationships between neuronal activity in V1 and SC and behavioral detection of visual contrast or luminance. The top panel of Figure 2A combines data from 8 PV+ mice during 209 contrast change detection sessions where white noise optogenetic inhibition was delivered to V1. The average NBK is comparable in breadth to that from the single session in Figure 1D, indicating that the timing of effects across animals and sessions is largely consistent. The negative peak of the average V1 Gabor NBK occurs 57 ms after the onset of the visual stimulus (FWHM = 51 ms; from 25 to 76 ms after visual stimulus onset). Activation of V1 PV+ neurons before the visual stimulus or 200 ms after visual stimulus onset had no obvious effect on performance, although the visual stimulus remained on the display until the end of the trial. The bottom panel of Figure 2A shows the combined NBK resulting from white noise optogenetic stimulation of GAD2+ neurons in SC for detection of visual contrast changes (7 mice, 185 sessions). The NBK resulting from SC stimulation was similar to the NBK obtained from V1, with the initial negative peak occurring 48 ms after the onset of the visual stimulus (FWHM = 77 ms; 37 to 114 ms). These perturbations suggest that spiking activity in SC and V1 are both causally linked to the behavioral response in this task and that the earliest moments of spiking in both brain areas contributes most to performance. The peaks of the V1 and SC NBK’s occur ∼250 ms before the peak in the reaction time distribution (see superimposed response time histograms). To quantify the reliability of these effects, we computed the area over the NBK (AOK) using a bootstrap procedure. For each of 100 bootstrapped NBKs, we sampled with replacement from the optogenetic stimulus profiles from trials that ended in hits and misses and computed the integral of the difference from 0 normalized power during the first 100 ms following the onset of the visual stimulus. Figure 2B shows that the distribution of bootstrapped AOKs was significantly different from 0 for both V1 and SC (V1 median = 1.29; SC median = 1.03; both p < 0.05).

**Figure 2.**
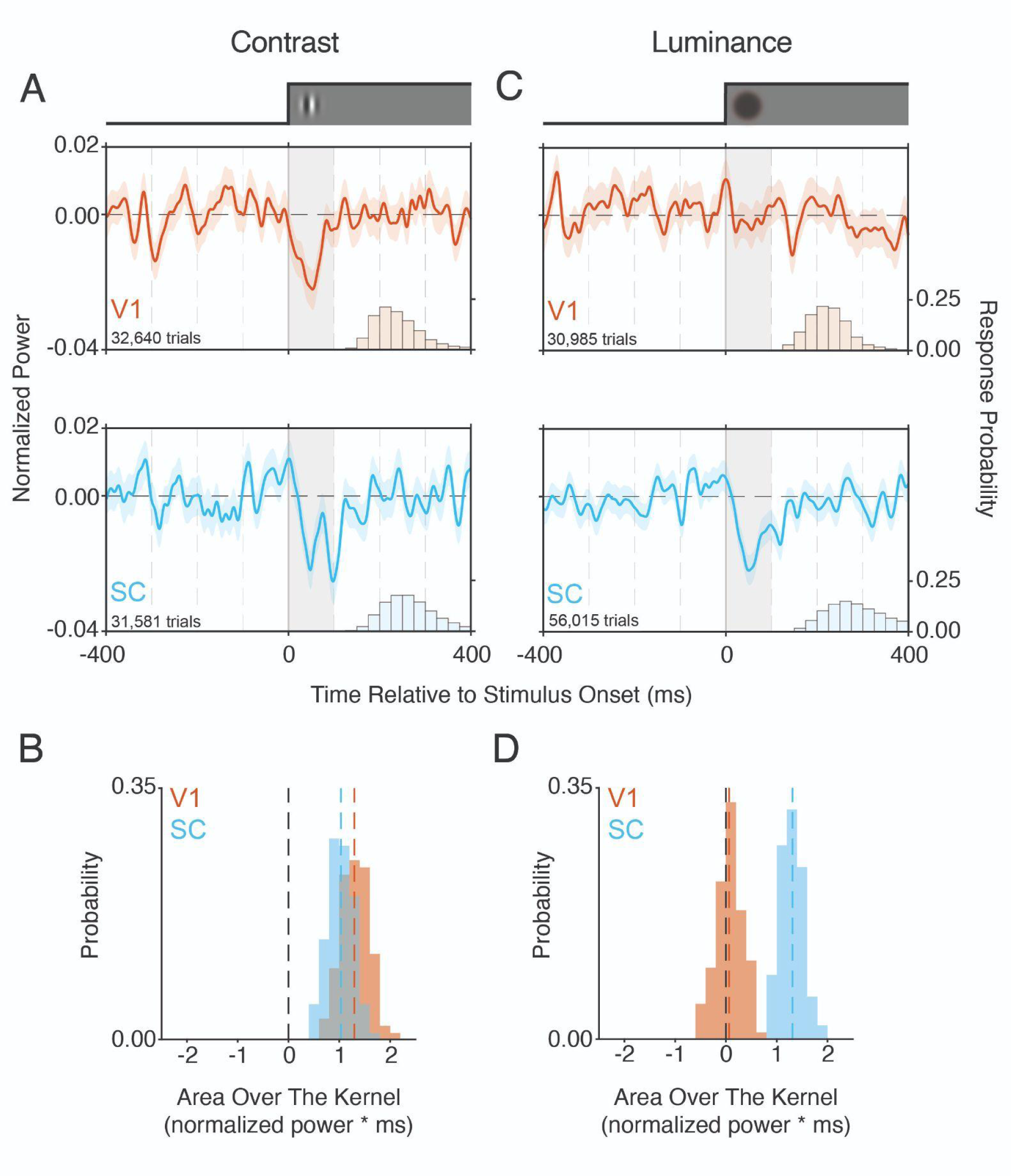
Neuronal-behavioral kernels (NBKs) from V1 and SC reveal stimulus dependent contributions to perception. **A)** (top) V1 NBK (red) aligned to the onset of visual contrast changes (t = 0) constructed by combining trials with optogenetic stimulation from 209 sessions from 8 mice. The superimposed histogram shows (right y-axis) a portion of the reaction time distribution. (bottom) The SC NBK (blue) aligned to visual contrast changes, combines trials with optogenetic stimulation from 185 sessions from 7 mice. Gray box from 0-100 ms highlights analysis window used in B). Shaded area represents ±1 SEM. **B)** Distribution of bootstrapped area over the NBK (AOK) from 0-100 ms following stimulus onset for SC (blue) and V1 (red). Both distributions are significantly different from zero (p<0.05, bootstrap). Colored dashed lines mark the medians of their respective distributions. **C)** Same as in A except the V1 NBK (top) combines data from 163 sessions in 6 mice and the SC NBK (bottom) combines data from 367 sessions in 11 mice during luminance change detection sessions. **D)** Same as in B except the AOK distributions were computed from luminance detection sessions. The V1 distribution is indistinguishable from zero suggesting that optogenetic stimulation during the 100 ms following visual stimulus onset has no systematic relationship with detection performance, while the SC distribution is significantly different from zero (p<0.05, bootstrap).

Figures 2C-D show the results from V1 (6 mice, 163 sessions) and SC (11 mice, 367 sessions) in which mice detected changes in visual luminance. Unlike the NBKs for detection of visual contrast changes, there were clear differences between the NBKs obtained from V1 and SC. Figure 2C (top) shows the V1 NBK for detection of luminance changes. Strikingly, there were no strong modulations in the V1 NBK following the onset of the visual stimulus. However, the SC NBK (Figure 2C, bottom) showed a strong modulation that peaked 51 ms following visual stimulus onset (FWHM = 49 ms; 25 to 74 ms). Similar to contrast detection sessions, the peak of the SC luminance NBK occurred ∼250 ms before the peak of the reaction time distribution. These observations were supported by the AOK bootstrapping procedure as the median AOK computed during the first 100 ms after visual stimulus onset for V1 was near 0, while the AOK distribution from SC was significantly different from 0 (Figure 2D; V1 median = 0.06, p = 0.37; SC median = 1.31, p < 0.05). We next asked whether the four AOK distributions originated from the same underlying distribution and found that not to be the case (p < 0.0001; Kruskal-Wallis test). Post-hoc tests revealed that the V1 Luminance AOK distribution was significantly different from all other distributions while the SC contrast AOK distribution was significantly different from the V1 contrast AOK and the SC luminance AOK distributions (all p < 0.0001, Dunn’s test). Figure S1 plots the AOKs obtained in individual sessions with the corresponding change in detection performance.

The striking difference in the effect of V1 inhibition with respect to detection of luminance versus contrast changes is intriguing. The average power delivered to V1 during luminance detection sessions was significantly greater compared to contrast detection sessions (see *Behavioral Performance* for mean powers across conditions; p < 0.001, Wilcoxon rank-sum test). We elevated the power for V1 luminance sessions in response to preliminary analyses indicating a lack of modulation in the V1 luminance NBK. Further, there were no differences in Δd’ for V1 luminance versus contrast sessions (see medians listed above; p > 0.05, Wilcoxon rank-sum test). Taken together with the NBKs, these data suggest that white noise inhibition of V1 during luminance detection sessions resulted in a small disruption in performance but the effects were not specific to any particular moment relative to stimulus onset. Consistent with this idea, we computed the AOK for V1 luminance sessions during the 400 ms before to 100 ms after visual stimulus onset and found the AOK distribution trended towards a difference from zero (median = −0.88, p = 0.07, bootstrap; note that due to our conventions, the negative median indicates that the AOK trends above 0).

In contrast, white noise inhibition of V1 during contrast detection sessions, even with lower mean powers, produced a temporarily specific effect on performance. Unlike V1, the mean power for SC stimulation during contrast detection sessions was greater than that used during luminance detection sessions (p < 0.0001, Wilcoxon rank-sum test). However, the effects of white noise stimulation on detection performance (Δd’) were greater for luminance detection sessions (p < 0.01, Wilcoxon rank-sum test) and in both cases the SC NBKs showed clear modulations time-locked to the earliest moments after stimulus onset. In sum, these data suggest that detection of luminance changes depends on the earliest periods of activity in SC but not V1, while detection of visual contrast relies on the earliest moments of activity in both structures.

The data in Figure 2 suggest a striking difference in the contributions of V1 spiking to detecting changes in contrast versus luminance. However, detection of some visual stimuli might be more susceptible to optogenetic perturbations than others. For example, detection of stimuli closer to the detection threshold are likely to be more impacted by an equivalent perturbation than stimuli near the extremes of a psychometric function. Furthermore, only a subset of V1 mice performed both contrast and luminance detection sessions (n=6). Thus, the effects shown in Figure 2 could depend on which mice contributed to the NBKs as well as differences in the effect of optogenetic perturbations on detection performance. We performed an additional analysis in which we only considered data from the subset of mice that performed both contrast and luminance detection sessions (136 sessions of each stimulus type). We further mean-matched the distributions of Δd’ for luminance and contrast change detection sessions so the respective NBKs would arise from equivalent effects of optogenetic perturbations on detection performance. The differences between the V1 NBKs for luminance and contrast were preserved by this more restrictive analysis (Figure S2) suggesting that the effects shown in Figure 2 were not carried by specific mice or by the magnitude of change in d’ induced by optogenetic stimulation.

An additional consideration is whether the effects reported above depend on the retinotopic alignment between the visual stimulus representation in SC or V1 and the patch of neurons perturbed by white noise optogenetic stimulation. While Figure 1B shows that the effects of optogenetic stimulation are spatially localized, the optogenetic stimulus in those experiments was a square pulse of fixed duration and intensity that was presented only during presentation of the visual stimulus. Stochastic optogenetic stimulation trains like those used here could produce deficits in detection performance that are independent of sensory input such as disrupting circuit function, distracting the animal, or interfering with motor preparation. We thus collected additional datasets from subsets of the same mice used above in V1 (n=5 mice, 122 sessions) or SC (n=6 mice, 156 sessions) in which the visual stimulus was moved at least 25° away from the optimally aligned location identified in Figure 1B. Migrating the visual stimulus away from the retinotopic location of white noise optogenetic stimulation eliminated the post-stimulus modulation in the NBK in both SC and V1 (Figure S3). These data argue that white noise optogenetic stimulation must be delivered to the neurons that represent the visual stimulus, arguing that the NBKs shown in Figure 2 reflect the moment-to-moment influence of optogenetic stimulation on behavioral detection rather than affecting performance through non-sensory mechanisms.

### Response-time-aligned kernels reveal distinct effects on behavioral responses

The NBKs in Figure 2 result from aligning optogenetic stimuli to the onset of visual stimuli (t = 0). This alignment is ideal for assessing the behavioral contributions of primary sensory areas like V1, however the SC could play a role in sensory decoding, motor preparation, or both. To address this, we computed response-time-aligned NBKs for V1 and the SC. To increase precision, we combined data from luminance and contrast detection sessions, since the motor response was the same for both. As before, we only considered sessions where Δd’ was greater than 1 SD of the mean. This yielded 355 sessions for V1 (n=8 mice) and 554 sessions from SC (n = 11 mice). First, we computed response-time-aligned NBKs for trials that ended in false alarms, where behavioral responses occur independent of the visual stimulus (Figure 3A). There was no obvious modulation in V1 false-alarm aligned NBK in the hundreds of milliseconds before lever responses (Figure 3A, top), suggesting that false alarms were not driven by fluctuations in V1 activity that might result in the animals perceiving a fictive visual stimulus (e.g., phosphene). It should be noted, however, that the smaller number of false alarm trials make detecting a signal more difficult. Nevertheless, this outcome is consistent with previous work showing that mice cannot detect isolated activation of their V1 PV+ neurons.^8,49^ In contrast, there was a noticeable negative deflection in the SC false-alarm aligned NBK occurring between 250 and 100 ms before spontaneous lever responses (Figure 3, bottom). This suggests that lever responses could be driven by momentary reductions in the optogenetic stimulus power ∼225 ms before lever responses. Figure 3B shows the bootstrapped distributions of the AOKs for V1 and SC during the 250 ms preceding false alarms. The V1 AOK distribution was not significantly different from zero (median = 0.14, p = 0.45), whereas the SC distribution was (median = 1.17, p = 0.02). Further, the false alarm aligned AOK distributions differed significantly (p< 0.0001, Kruskal-Wallis test), confirming the visual impressions from the two NBKs.

**Figure 3.**
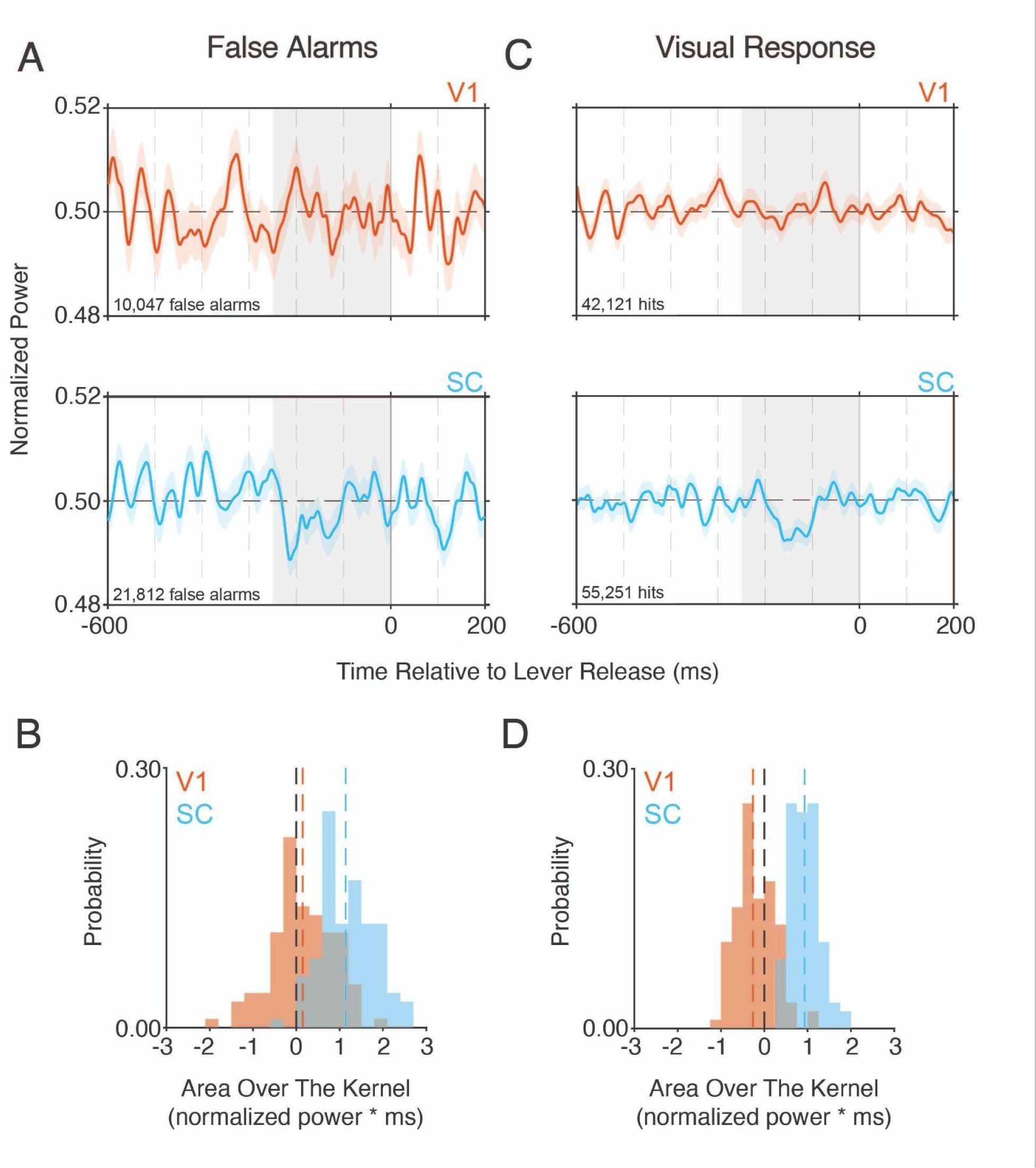
Neuronal-behavioral kernels aligned to behavioral responses. Data combines observations from contrast and luminance sessions for V1 (n=8 mice, 355 sessions) and SC (n=11 mice, 554 sessions). **A)** (top) V1 NBK (red) aligned to the onset of lever responses (t = 0) on trials where the mice responded prior to the onset of the visual stimulus (false alarms). (bottom) The SC NBK aligned to false alarms combining trials with optogenetic stimulation. Gray box from −250-0 ms highlights analysis window used in B. Shaded area represents ±1 SEM. **B)** Distribution of bootstrapped area over the NBK (AOK) from −250-0 ms following stimulus onset for SC (blue) and V1 (red). Colored dashed lines indicate the medians of their respective distributions. Only the SC false alarm AOK distribution is significantly different from zero (p<0.05, bootstrap) and the two AOK distributions are significantly different from one another (p<0.0001, Kruskal-Wallis test). This shows that false alarms were more likely when the SC optogenetic stimulus power was lower than average during this epoch while changes in V1 activity is not strongly associated with producing false alarms. The negative peak in the SC NBK occurs ∼225 ms prior to lever releases suggesting that release from SC inhibition during this window may generate a fictive percept (e.g. phosphene) that mice are using to guide their responses. There was no obvious modulation in the V1 false alarm NBK supporting our prior work showing that inhibition of V1 cannot support perception.^8,49^ **C)** Same as in A except the V1 (top) and SC (bottom) NBKs are aligned to lever releases on hit trials. Similar to the V1 false alarm NBK, there is no obvious modulation in the V1 hit NBK at any time prior to lever responses. The SC hit NBK exhibits a negative modulation likely due to broadening of the NBK that is time-locked to stimulus-onset. **D)** Same as in B except the AOK distributions were computed from hit trials. The V1 distribution does not differ from zero (p>0.05) while the SC distribution is significantly different from zero (p<0.05, bootstrap) while the two AOK distributions are significantly different from one another (p<0.0001, Kruskal-Wallis test). Comparing neuronal behavioral kernels with neuronal activity in V1 and SC

Next, we examined response-time-aligned NBKs on hit trials, where behavioral responses occur in response to the presentation of visual stimuli (Figure 3C-D). The NBKs computed from hit trials in V1 and SC were similar to their respective false-alarm aligned kernels. There were no signs of a strong modulation in the response-time-aligned V1 NBK (Figure 3C, top) but a clear negative deflection in the SC response-time-aligned NBK (Figure 3C, bottom). The modulation in the SC NBK was broader than the visual stimulus-aligned NBKs in Figure 2, suggesting that the spikes in SC that drive behavioral detection of a visual stimulus are those that occur time locked to the onset of that stimulus. The lack of a comparable modulation in the response-time-aligned V1 NBK likely stems from two factors: 1) we combined data from both luminance and contrast detection sessions and there were no obvious modulations in the visual stimulus-aligned NBK for luminance in V1 (see Figure S4 for response-time-aligned NBKs for the individual visual stimulus conditions) and 2) the stimulus aligned NBKs from V1 separated for hit and miss trials shown in Figure S5 show a stronger modulation in the optogenetic stimulus power for misses. Together these two features would strongly dilute the stimulus related signal in the response-aligned V1 NBK. The distributions of bootstrapped AOKs corroborated our results (V1 AOK median = −0.6, p > 0.05; SC AOK median = 0.93, p < 0.05). Moreover, the response-time-aligned AOK distributions were significantly different from one another (p< 0.0001, Kruskal-Wallis test). Importantly, the temporally restricted effects associated with sustained white noise optogenetic stimulation rule out behavioral effects arising from the optogenetic stimulus causing cortical heating, direct retina stimulation, or nonspecific behavioral effects. Furthermore, it is possible that removing inhibitory input in SC independent of a visual stimulus can produce signals that mice can use to guide their behavioral responses but directly addressing this is beyond the scope of this study.

The early peaks in the V1 and SC NBKs suggest that behavioral detection is preferentially driven by the earliest spikes in the response to a visual stimulus. However, the effects of optogenetic stimulation on spiking are delayed relative to the optogenetic stimulus because of the kinetics of the opsin as well as synaptic delays between inhibitory neurons and their synaptic partners.^43,50^ Thus, we used extracellular electrophysiology to examine how the optogenetic stimulus impacted neuronal spiking in V1 and SC. The relevant delays can be captured by recording neuronal responses and computing the spike triggered average (STA) in response to white noise optogenetic stimuli. The STA represents the impulse response of a spiking neuron (Bryant and Segundo, 1976). The aggregate STA across a neuronal population provides the overall neuronal response to an impulse of optogenetic stimulation, capturing the relevant delays to changes in spiking across the population.

We computed the STAs by performing a reverse correlation of the optogenetic stimuli aligned to the occurrence of individual spikes (at t = 0). We measured significant STAs in 29/56 units recorded in V1 and 22/105 units in SC. Figure 4 depicts the population STAs from V1 and SC which characterize the average delays between impulses of optogenetic stimulation and changes in spiking. Figure 4A (top) shows the STA of a single V1 unit that was excited by optogenetic stimulation (putative PV+ cell expressing ChR2), as indicated by the large positive peak in the optogenetic power immediately before the occurrence of spikes (∼1 ms). The STA closely resembles previously published observations from V1 (Day-Cooney, Cone, and Maunsell 2022), supporting the idea that putative PV+ cells respond with short latency following a step in the optogenetic stimulus power. Note that the peak of the STA is artificially widened by pulse width of the white noise optogenetic stimuli (25 ms) leading the positive peak to extend beyond the time of spikes (t=0), where optogenetic stimulation could not possibly influence spiking. The combined STAs for 2 SC units that were excited by optogenetic stimuli are shown in the bottom panel of Figure 4A (e.g., putative Gad2+ neurons expressing ChR2). Much like V1, spikes occur almost immediately (∼1 ms) following the positive peak in the optogenetic stimulus power.

**Figure 4.**
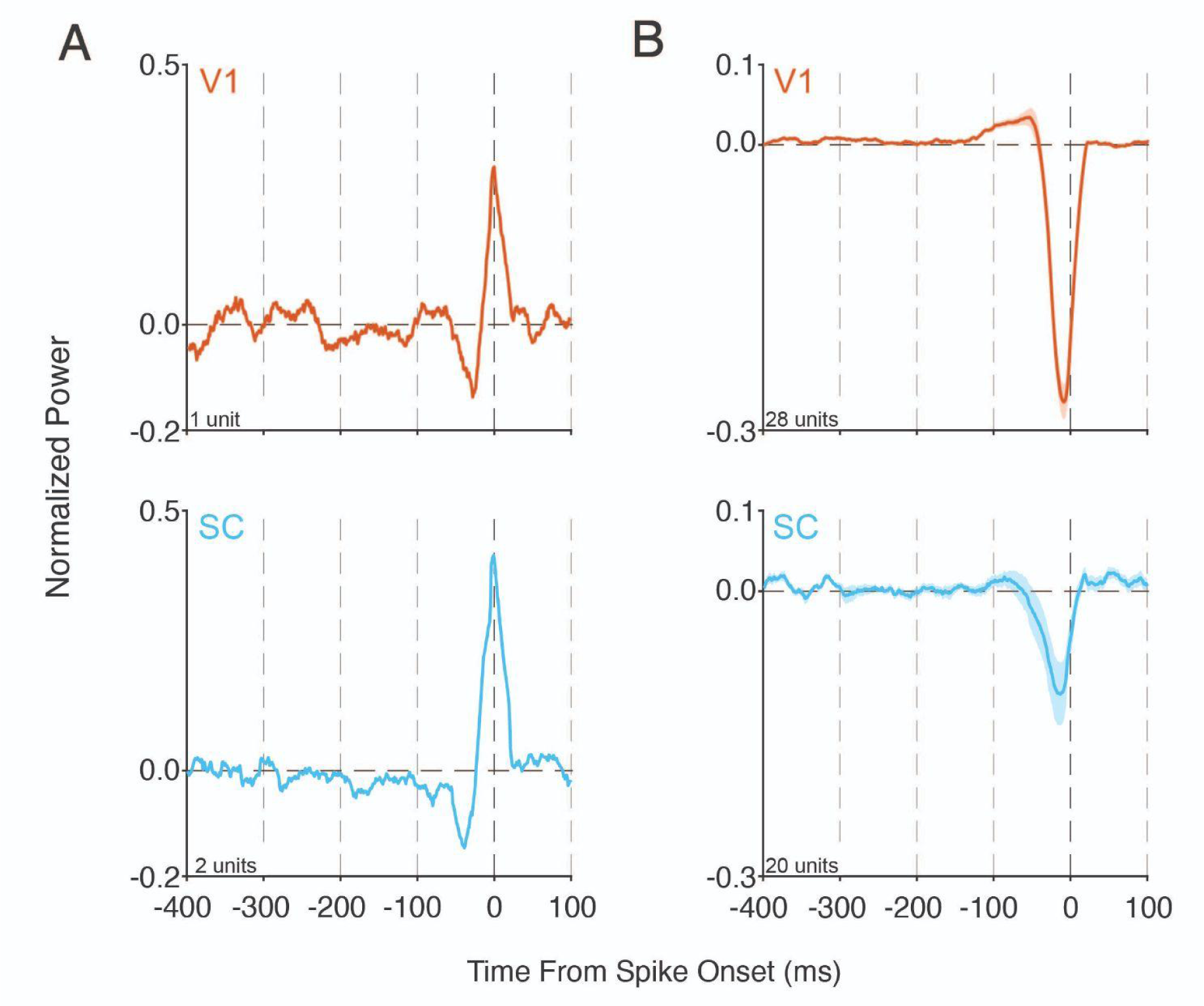
Spike-triggered average optogenetic stimuli reveal the time course of white noise optogenetic stimulation effects on spiking. Optogenetic stimulus profiles aligned to the time of V1 (red) or SC (blue) spikes (t=0) were averaged to calculate a population STA optogenetic stimuli. A) (top) Red trace depicts the STA from one putative PV+ unit in V1. The unit spikes immediately (∼1 ms) after positive changes in the optogenetic stimulus. (bottom) Same as above except the blue trace represents the average STA 2 SC units that were excited by optogenetic stimuli. B) (top) The remaining V1 units (red trace) were inhibited by optogenetic stimulation as spikes tend to occur ∼10 ms following reductions in the optogenetic stimulus power. (bottom) Same as above except for the remaining SC units that were inhibited by optogenetic stimulation. Shaded area in B represents ± 1 SEM.

The average STA for V1 units that were not directly excited by optogenetic stimulation (e.g., putative principal neurons that receive input from PV+ cells, n = 28) is plotted in Figure 4B. The V1 STA (top) shows that on average these V1 neurons tend to spike ∼10 ms following decreases in the optogenetic stimulus power. There is a slight positive modulation in the optogenetic power that peaks about 50 ms before spikes, indicating that spikes are more likely if PV+ stimulation power is higher than the average during this period. However, the magnitude of the positive modulation is less than 20% of the dominant negative peak immediately preceding spikes. Figure 4B (bottom) shows the corresponding results for 20 SC units where spikes also tended to occur ∼10 ms following decreases in the optogenetic stimulus power. Overall the timing of the negative peak in the STA and the dynamics of the optogenetic power surrounding spikes in SC and V1 were strikingly similar. Taken together, the data in Figure 4 show that the dominant effects of white noise optogenetic stimulation on V1 and SC spiking, and thus the NBK, lag the optogenetic stimulus by ∼10 ms and are largely restricted to the moments immediately preceding spikes.

An important question is the extent to which neuronal responses in V1 and SC encode information about the visual stimuli that we used to estimate the NBKs. For example, the absence of a modulation in V1 luminance NBK suggests that mice do not appreciably weight V1 spikes when detecting changes in luminance. However, a simple explanation for this result could be that mouse V1 neurons do not respond strongly to changes in luminance and therefore do not carry information that mice could use to perform the task. To address this, we also did acute electrophysiological recordings in which we presented luminance or contrast changes in a subset of mice we used to estimate the STAs (V1: 9 mice, 16 recording locations; SC: 7 mice, 14 recording locations). Visual stimuli were positioned inside the receptive field of a randomly selected unit on the electrode array (see Materials and Methods). Visual stimuli were presented on every trial, with a random half of trials including white noise optogenetic stimuli so we could compute the trial-averaged effect of optogenetic stimulation on the respective populations.

We encountered units that responded to both stimulus types in SC and V1 and the majority of these units were excited by visual input (Figure S6). Overlaying the visual responses with the FWHM of the NBKs suggests that the earliest spikes in SC and V1 contribute preferentially to stimulus detection. We next sought to assess the amount of information carried by V1 or SC populations for contrast and luminance. We did an ROC analysis of the spike counts from the subpopulations of V1 or SC neurons that were excited by contrast or luminance changes. The ROC tested the ability to discriminate the onset of a visual stimulus during the first 100 ms after visual stimulus onset from the baseline spike counts on each trial. This analysis revealed that both V1 and SC units carried substantial information about the onset of visual stimuli (auROC: V1 contrast: 0.87; SC contrast: 0.71; V1 luminance: 0.79; SC luminance: 0.74; Figure S7). These data show that V1 and SC populations both convey information that can be used to detect changes in luminance. Despite the potential information carried by V1 units for luminance stimuli, the NBK shown in Figure 2 argues that mice do not use these signals to guide their behavioral responses.

Next, we determined the average change in spiking produced by white noise optogenetic inhibition on the visual responses quantified above. We computed the average firing rate of each recorded unit for trials with versus without optogenetic stimulation. Because the presence/absence of optogenetic stimulation was randomized across time bins, we used the average firing rate computed from the entire trial (50 ms bins). Trials without optogenetic stimulation were the same as those used to analyze visual responsiveness (see above). The firing rate of many neurons in V1 and SC were significantly modulated by optogenetic stimulation (V1 contrast: 60/93 units; SC contrast: 32/78 units; V1 luminance: 54/113 units; SC luminance: 40/112 units; p < 0.05, Wilcoxon signed-rank test,). As expected, the vast majority of modulated units were inhibited by optogenetic stimulation regardless of brain area or stimulus type (V1 contrast: 54/60; SC contrast: 27/32; V1 luminance: 41/54; SC luminance: 34/40). In keeping with the modest changes in behavioral detection performance (i.e., Δd’), the overall impact of white noise optogenetic stimulation on the trial average spike rate was small but significant (mean Δspikes/s, V1 contrast: −0.99, SEM: 0.26; SC contrast: −0.24, SEM: 0.38; V1 luminance: −0.69, SEM: 0.39; SC luminance: −0.33, SEM: 0.24; difference from 0 all p<0.005, Wilcoxon signed-rank test). Thus, on average, the powers chosen for white noise stimulation resulted in small reductions in spiking in both SC and V1 that were consistent across stimulus types.

### Electrophysiological Interactions

Complex interactions between visual processing in V1 and SC have long been appreciated.^51^ SC receives strong input from V1.^52,53^ Likewise, SC projects to both the lateral geniculate (LGN) and the lateral posterior (LP) nuclei of the thalamus.^54^ Recent work has demonstrated that inhibition of the SC can reduce V1 visual responses.^55^ Similarly, inhibition of V1 reduces the response of SC neurons to visual input.^56^ Thus, to assess whether white noise optogenetic perturbations of SC disrupted processing in V1 (and vice versa), we performed electrophysiological recordings in V1 or SC while we delivered white noise optogenetic stimulation to the other brain region. The recording electrode was positioned in the same receptive field location as the optogenetic perturbations. We recorded from 37 units in V1 (n=3 mice, all male) while delivering white noise optogenetic stimulation to opsin expressing GAD2+ SC neurons. We then aligned SC optogenetic stimuli with V1 spike times to compute a cross area STA. There were no obvious modulations in the SC optogenetic power at any time before V1 spikes, suggesting that the white noise optogenetic stimulation in SC did not appreciably augment V1 spiking (Figure S8A). We also recorded from 17 SC units (n=2 mice, 1 female) while delivering white noise optogenetic stimulation to opsin expressing V1 PV+ neurons and computed a cross-area STA. There were similarly no detectable modulations in the V1 optogenetic stimulus power before SC spikes (Figure S8B). Together, these data support the idea that while V1 and SC share reciprocal direct and indirect connectivity, white noise optogenetic stimulation delivered to V1 or SC at the powers used during behavioral sessions did not appreciably disrupt spiking in the other structure. Lastly, we also asked whether any of the effects we observed resulted from non-specific disruption of neuronal activity independent of opsin activity (i.e., tissue heating). We recorded from 15 V1 units in 2 opsin naive mice (1 female) while delivering white noise illumination via the optical fiber. There were no detectable modulations in the V1 population STA at any time proceeding spikes (Figure S8C). These data suggest that modulations in the STA (and NBK) result from direct activation of opsin expressing neurons and not from indirect changes in activity due to tissue heating.

## Discussion

Reverse correlation is a powerful tool that has led to important insights into the dynamics of many neurobiological processes including receptive field properties,^46,57^ perceptual decision making,^58,59^ sensory processing,^60^ and visual attention.^61^ Here, we used white noise optogenetic stimulation of inhibitory neurons in V1 or SC while mice engaged in a challenging visual detection task, leading to three primary observations. First, SC and V1 both contribute to behavioral detection of changes in visual contrast. Second, behavioral detection of changes in visual luminance depends on the SC but not V1. Lastly, the contributions of SC and V1 to behavioral detection of visual stimuli are limited to the first 100 ms after the onset of visual stimuli.

The presence and absence of a detectable V1 NBK during contrast and luminance detection, respectively, underscore the complications associated with evaluating perturbations of neuronal circuits. We interpret the presence of a NBK during contrast detection as indicating that the spiking of V1 neurons contributed directly to the detection of contrast targets. The absence of a NBK during luminance detection points instead to V1 perturbations generating some other change in behavioral state that acted to distract the animal from the task. Such effects might have persisting impacts on performance, and therefore the optogenetic signals from different trials would not combine to produce a NBK restricted to a particular interval relative to stimulus onset or lever release.

If optogenetic stimulation of V1 impacted performance in this way, similar effects might also have existed during contrast detection sessions. However, this effect could coexist with a critical contribution of V1 spiking to detecting contrast stimuli, and would not necessarily mask the kernel associated with that contribution. It is conceivable that the similar small decreases in d’ associated with optogenetic stimulation in both tasks (median Δd’ for contrast −0.06, for luminance −0.05) depended entirely on such effects. Because white noise optogenetic stimulation increases the probability of success on some trials and decreases that probability on others (Figure 1D), the presence or absence of a temporally restricted NBK is not necessarily linked to any difference in d’. Without the NBK analysis, we would have simply observed reduced performance on both contrast and luminance detection during V1 inhibition and inferred that V1 activity was involved in both tasks - a valid, but imprecise assessment. Weak, white noise stimulation with NBK analysis allowed us to resolve its strikingly different contributions to these outwardly similar behaviors.

While the SC has long been appreciated for its role in overt and covert orienting behaviors,^10,16^ its prominent NBK reinforces a growing list of studies suggesting a role in visual perception.^17,25,26,30^ Approximately 80% of RGCs in mice project to both the LGN and the SC, implying that similar information is represented in SC and V1.^27^ Indeed, many visual features traditionally associated with cortical visual processing are also encoded in the responses of SC neurons.^19–24^ While the role of V1 in contrast perception is well established,^8,47,62^ the distribution of neuronal contrast sensitivities are similar between V1 and the SC in the rat^63^ making the SC well-positioned to provide contrast related signals.

The lack of a V1 contribution to luminance detection was unexpected, especially in light of the robust V1 responses to changes in luminance observed by us as well as by others.^64,65^ While sensory inputs activate neurons throughout the brain,^66,67^ subjects preferentially weight the signals that are most relevant for perceptual decisions.^68–71^ The SC is capable of generating perception independent of V1. Human observers with V1 lesions can exhibit paradoxically lower contrast detection thresholds in the affected visual field representation compared to the intact portions.^72^ The ability to sense and respond to visual inputs in parts of the visual field affected by damage to V1 (e.g., “blindsight”) is believed to be due to contributions from the SC.^5^ Similarly, the Sprague effect, in which loss of visual function following unilateral ablation of V1 can be improved by contralesional ablation of the SC, shows that the SC ipsilateral to the V1 lesion can mediate aspects of vision that are traditionally associated with visual cortex.^51^ The absence of a detectable V1 NBK during luminance detection points to V1 responses being less useful for responding to luminance stimuli. Activating inhibitory neurons before or after the earliest portion of V1 or SC responses had no detectable effect on behavioral performance, a result that is supported by both experimental and modeling work.^25,31,73^ This result, together with the absence of a V1 NBK during luminance detection, underscores the potential pitfalls of assuming that neuronal signals conveying task-relevant information are used to guide behavior: V1 neurons were activated by visual luminance but there was no corresponding modulation evident in the V1 luminance NBK. It is possible that performance would have improved were animals to combine luminance information from SC and V1. While subjects can optimally integrate distinct sensory signals for guiding behavior, ^68,74^ there are also many examples where subjects clearly fail to do so.^49,75–78^ Whether such failures arise from suboptimal behavioral strategies or neurobiological limitations the decoding of sensory information from neuronal populations warrants future investigation.

Aligning the optogenetic stimulus profiles to behavioral responses instead of visual stimulus onset also revealed differences between V1 and SC. The behavioral response-aligned NBKs largely conform to traditional views of SC and V1 where the SC subserves sensorimotor functions while V1 is predominantly a sensory area. There were no obvious modulations in the V1 NBK aligned to false alarms, which supports the idea that direct inhibition or release from inhibition in V1 fails to generate signals that mice can use to guide their behavior.^49^ In contrast, the SC false alarm aligned NBK shows a modulation that precedes lever releases by about 200 ms. The long delay from the NBK modulation to the lever response is unlikely due to a direct drive of forelimb movements but is most likely to result if release from SC inhibition generates a weak fictive visual stimulus (e.g., phosphene) that the mouse uses to guide its responses. If so, this suggests a striking asymmetry in the readout of signals from SC versus V1 because we have previously shown that mice do not respond to reductions in V1 spiking, nor to the increments in activity that follow a release from inhibition.^8,49^

The electrophysiological data support a temporally and spatially restricted effect of white noise inhibition on their respective circuits. While the population STAs from SC and V1 differ in magnitude, the time course of the effects of optogenetic stimulation on spiking are strikingly similar. Magnitude differences could arise for multiple reasons such as non-uniform ChR2 expression or the distance of the recorded neuron from the optical fiber. However, the comparable dynamics were unexpected in light of the substantially different molecular, circuit, and synaptic architecture in SC and V1.^32,54,79–81^ Cortical PV+ neurons provide rapid inhibition, primarily synapse on or near the soma, and are capable of firing hundreds of spikes per second.^81^ Gad2+ neurons in the SC spike at low rates and encompass both local interneurons as well as long-range projection neurons.^80,82^ Despite these differences, the STAs suggest similar time courses for changes in spiking following pulses of optogenetic stimulation. Moreover, we didn’t observe any consistent patterns in the optogenetic stimulus delivered to SC (V1) when aligned to V1 (SC) spikes. This suggests that at the powers used in our study, the effects of optogenetic stimulation on spiking were not mediated by effects of one structure on the other. While others have shown that optogenetic inhibition of SC (V1) can modulate visual responses in V1 (SC), these differences are most likely because we used powers that were orders of magnitude lower than prior work.^55,56^

These data show that white noise optogenetic stimulation can be successfully deployed in brain areas with vastly different developmental, genetic, and circuit architectures. Importantly, we can recover NBKs even at low mean optogenetic stimulus powers that produce only modest effects on behavior (as measured by Δd’). A key advantage of this approach is that keeping the optogenetic stimulus power low enables circuits to operate close to their normal operating ranges. While using higher optogenetic stimulus powers might yield NBKs in fewer trials, the possibility of inducing artifactual effects on circuit dynamics (e.g., long-lasting oscillations) are higher, which would likely contaminate temporal estimates of the contributions of spiking activity to behavior. Brain areas outside of the cerebral cortex, including the SC, feature greater cell-type diversity, but this could be overcome by directing opsin expression with promoter specific viruses. Furthermore, using opsins with faster off-kinetics such as Chronos could potentially yield NBKs with greater temporal precision.^83^ Although we collected NBKs from different brain areas in different mice, simultaneous white noise stimulation of multiple brain areas could yield multiple concurrent NBKs collected in the same trials and potentially highlight interdependencies between areas.

Despite these strengths, one limitation of this study is that we did not simultaneously perturb SC and V1 activity in the same animals. While PV+ neurons are ideally suited for cortical inhibition, PV is not a reliable marker of SC inhibitory neurons.^79^ We targeted SC inhibitory neurons using Gad2-cre mice as others have used this strain to study the role of SC in sensory decision making.^34^ In preliminary experiments (not shown), we found that V1 NBKs derived from activation of Gad2+ neurons were considerably noisier than our prior work using PV+ interneurons.^31^ This likely stems from the complex mixture of circuit effects that would result from activating the diversity of cortical interneurons labeled by Gad2.^33^ In particular, activation of cortical VIP+ inhibitory neurons can produce a behaviorally detectable signal.^8^ Another challenge is that NBKs are specific to a given stimulus and task parameters.^31^ Estimating an NBK requires many trials to repeatedly sample all periods surrounding stimulus onset. Here, we tested a single stimulus intensity for luminance and contrast that was below saturation but above threshold. It is possible that mice would leverage both V1 and SC activity if luminance stimuli were presented at perceptual threshold. Nevertheless, the data point to a striking asymmetry with respect to how SC and V1 signals contribute to detection of changes in luminance.

## Acknowledgements

The authors would like to thank Dr. Patrick Mayo, Chery Cherian, and members of the Cone lab for helpful feedback on the manuscript. This work was supported by NIH grants R21-EY032650 (J.J.C. and J.H.R.M.) and U19-NS107464 (J.H.R.M.) as well as an NSERC Discovery grant RGPIN-2022-03283 (J.J.C.).

## Supplemental Information

**Supplemental Tables.** Behavioral performance by visual and optogenetic stimulus conditions. Tables report the median and interquartile intervals (IQR, in brackets) for d’ and c. * indicates the distribution was significantly different from zero (at least p < 0.05, Wilcoxon signed-rank test).

| 50% contrast | V1 contrast | SC contrast | V1 luminance | SC luminance |
| --- | --- | --- | --- | --- |
| $d'$ | 2.79 [2.4 3.1] | 2.44 [2.2 2.8] | 2.95 [2.6 3.4] | 2.56 [2.1 3.0] |
| $c$ | 0.66 [0.4 0.8] | 0.50 [0.3 0.7] | 0.59 [0.4 0.8] | 0.47 [0.3 0.6] |

| 30% contrast | V1 contrast | SC contrast | V1 luminance | SC luminance |
| --- | --- | --- | --- | --- |
| $d'$ | 2.53 [2.2 2.9] | 2.16 [1.8 2.5] | 2.70 [2.4 3.1] | 2.23 [1.8 2.7] |
| $c$ | 0.74 [0.6 1.0] | 0.65 [0.5 0.8] | 0.72 [0.6 0.9] | 0.64 [0.4 0.8] |

w/ white noise optogenetic stimulation
| 30% contrast | V1 contrast | SC contrast | V1 luminance | SC luminance |
| --- | --- | --- | --- | --- |
| $d'$ | 2.49 [2.1 2.8] | 2.07 [1.7 2.5] | 2.65 [2.4 3.1] | 2.14 [1.6 2.6] |
| $c$ | 0.77 [0.6 1.0] | 0.69 [0.5 0.9] | 0.75 [0.6 0.9] | 0.66 [0.5 0.9] |

Change in performance for 30% contrast with - without white noise optogenetic stimulation
|  | V1 contrast | SC contrast | V1 luminance | SC luminance |
| --- | --- | --- | --- | --- |
| $\Delta d'$ | -0.05 [-0.2 0.1]* | -0.05 [-0.2 0.1]* | -0.06 [-0.2 0.0]* | -0.08 [-0.2 0.0]* |
| $\Delta c$ | 0.03 [0.0 0.1]* | 0.02 [0.0 0.1]* | 0.03 [0.0 0.1]* | 0.04 [0.0 0.1]* |

**Figure S1.**
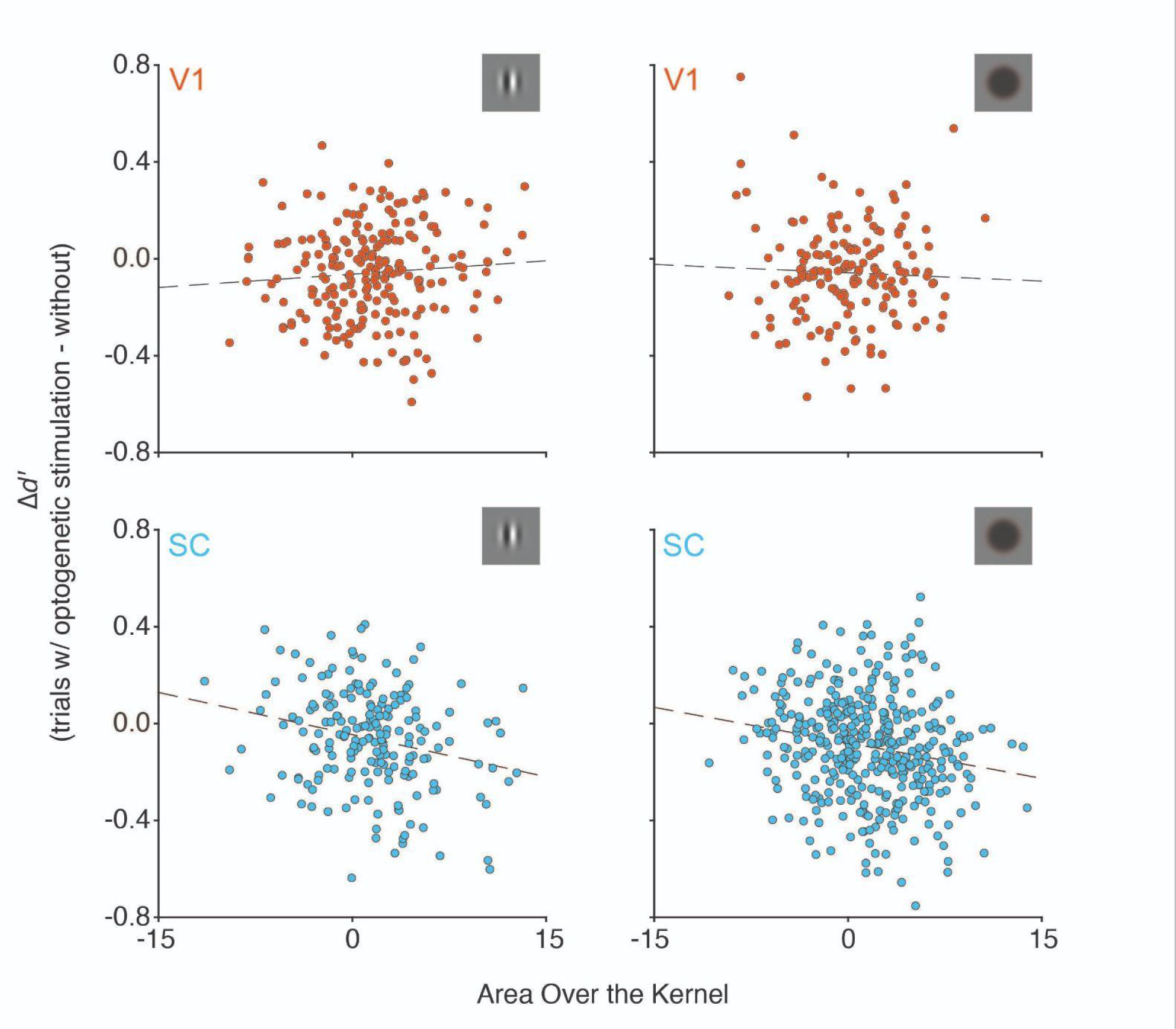
Scatterplots of the area over the NBK (AOK) by Δd’ for all individual behavioral sessions. The value for the AOK is the negative integral of the single session NBK. Positive values correspond to a larger negative deflection in the NBK like those shown in Figures 1D and 2A. The Δd’ is the difference between d’ on trials with optogenetic stimulation versus without. Negative Δd’ values correspond to larger performance deficits in the corresponding session. Each point (V1, red; SC, blue) represents values obtained in an individual behavioral session (all sessions shown). Gray box in the upper right corner indicates the visual stimulus type. Dashed lines represent the best fit line via simple linear regression. Given the stochastic nature of the optogenetic stimulus and the relatively low average powers used, not all sessions produce a reliable NBK. Thus, the AOK obtained in individual sessions was only weakly related to the impact of optogenetic stimulation on detection performance (correlation: V1 contrast (top left): ϱ = 0.08; p = 0.23; SC contrast (bottom left): ϱ = −0.24; p < 0.001; V1 luminance (top right): ϱ = −0.044; p = 0.58; SC luminance: ϱ = −0.2; p < 0.001).

**Figure S2.**
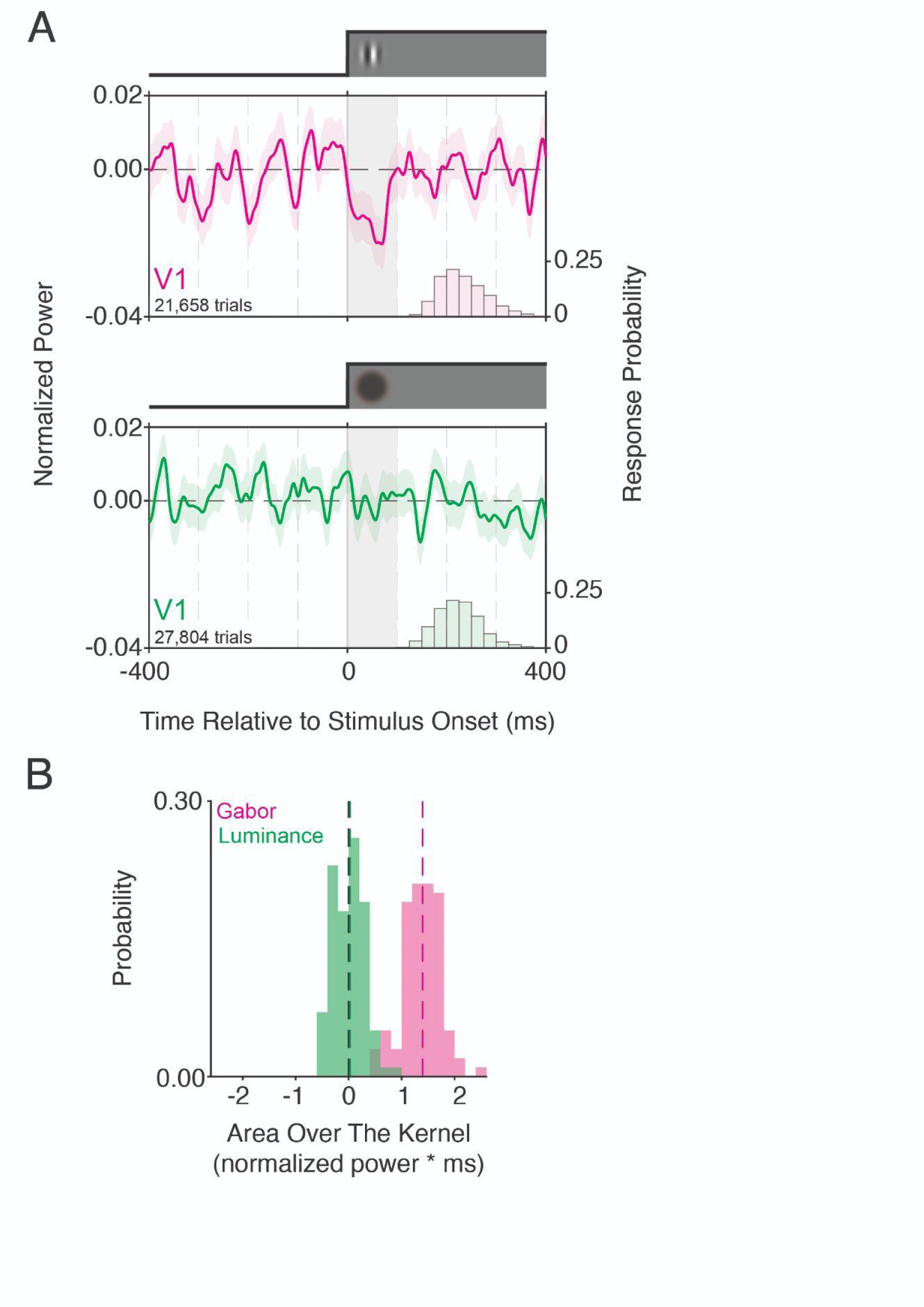
Animal and mean matched V1 NBKs for contrast or luminance changes computed from the same mice. Analyses were restricted to the 6 V1 mice who performed both contrast and luminance change detection sessions. 136 sessions of each stimulus type were included after mean-matching sessions for the effects of optogenetic stimulation on performance (mean Δd: contrast = −0.06, IQR = −0.17 - 0.04; luminance = −0.07, IQR = −0.17 - 0.04;). **A)** (top) The V1 NBK for visual contrast (pink) aligned to the onset of visual contrast changes (t = 0) constructed by combining trials with optogenetic stimulation. The superimposed histogram (right y-axis) shows a portion of the reaction time distribution. The V1 NBK for visual contrast exhibited a strong negative modulation during the initial 100 ms after visual stimulus onset, indicating that reduced V1 inhibition during this window is critical for detection performance. The profile of the visual stimulus is depicted directly above the plot. The analysis window used in B is indicated by the gray box. Shaded area depicts ±1 SEM. (bottom) Same as above except for sessions in which the same animals detected changes in luminance (green). There are no obvious modulations in the mean-matched V1 luminance NBK, suggesting that interfering with V1 spiking in the hundreds of milliseconds after the onset of luminance changes has no systematic effect on detecting that stimulus. The difference in trial counts between the two conditions is because mice tended to work longer during luminance detection sessions. **B)** Bootstrapped distributions of the area over the NBK (AOK) for contrast or luminance detection sessions computed from the data in A from 0-100 ms after stimulus onset. The distribution of AOKs for contrast was significantly different from zero (median AOK = 1.24, p < 0.05) while the distribution of AOKs for luminance was not (median AOK = 0.14, p = 0.32, bootstrap). Further, the AOK distributions for contrast versus luminance were significantly different from one another (p < 0.0001, Kruskal-Wallis test). The mean Δd and animal matched V1 NBKs for luminance and contrast closely resembled the results shown in Figure 2. This suggests that the differences between the V1 contrast and luminance NBKs do not depend on which mice contributed nor on the magnitude of impairment in detection performance produced by white noise optogenetic stimulation.

**Figure S3.**
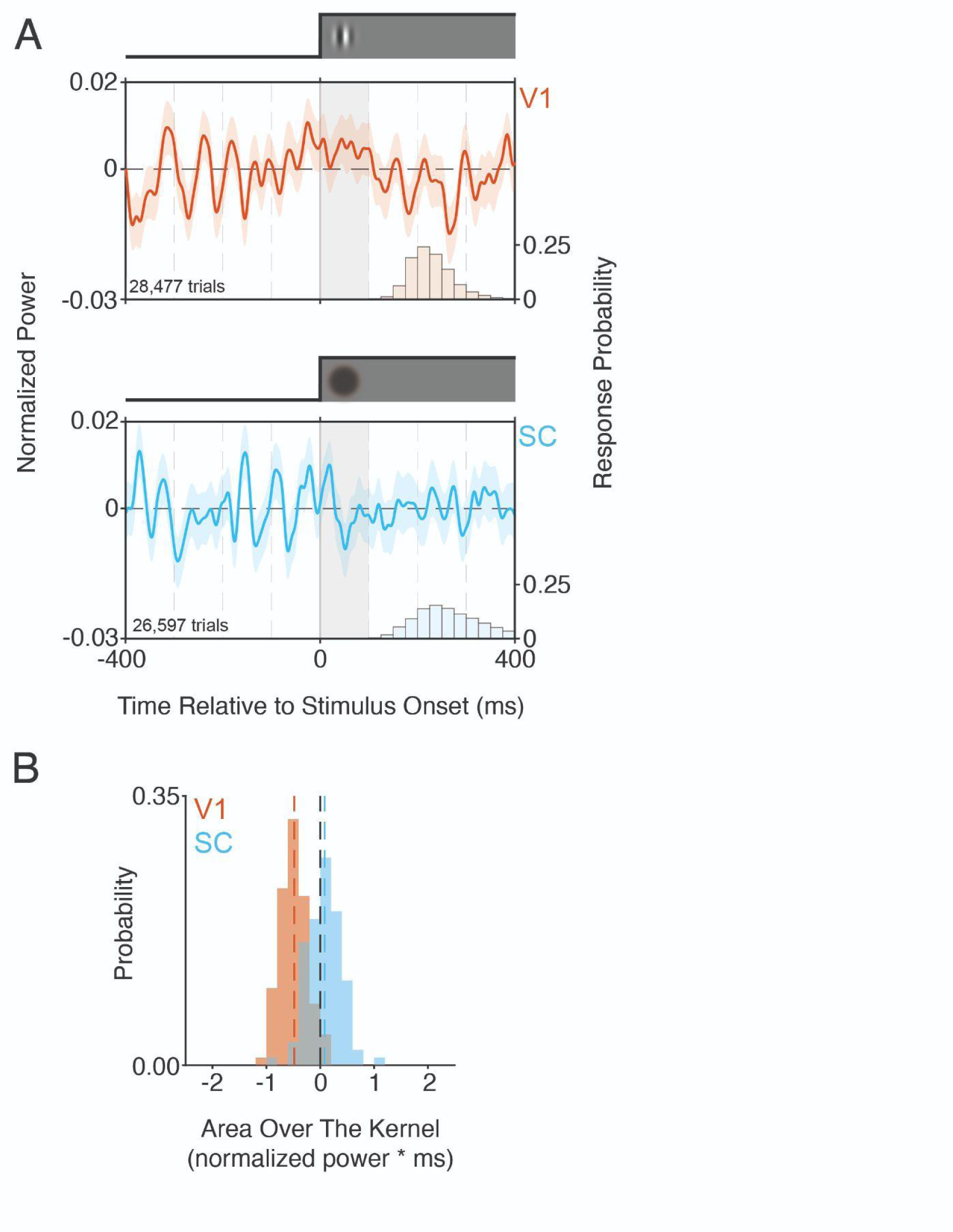
The effects of white noise optogenetic stimulation on detection performance depend on retinotopic alignment between visual and optogenetic stimuli. These data were collected from subsets of the same mice used in the main experiments (V1: n=5 mice, 122 sessions, median 23 sessions, range 19-39; SC: n=6 mice, 156 sessions, median 23 sessions, range 16-38). During these sessions, the visual stimulus was moved at least 25° away from the optimal location identified during preliminary alignment experiments (see Figure 1B). For this analysis, all sessions were considered regardless of the resulting Δd’. **A)** (top) The V1 NBK for visual contrast (red) aligned to stimulus onset (t = 0) constructed by combining trials with optogenetic stimulation. The superimposed histogram (right y-axis) shows a portion of the reaction time distribution. Migrating the visual stimulus away from the retinotopic location of optogenetic perturbations eliminates the negative modulation in the V1 contrast NBK seen in Figure 2A. The profile of the visual stimulus is depicted directly above the plot. The analysis window used in B is indicated by the gray box. Shaded area depicts ±1 SEM. (bottom) Same as above except for SC stimulation sessions in which mice detected changes in luminance (blue). Much like V1, moving the visual stimulus away from the site of optogenetic stimulation eliminates the negative modulation in SC luminance NBK shown in Figure 2C. **B)** Bootstrapped distributions of the area over the NBK (AOK) for contrast (V1) or luminance (SC) detection sessions computed from the data in A from 0-100 ms after stimulus onset. The distribution of the V1 AOK for contrast was significantly different from zero, although in the opposite direction as was observed when visual and optogenetic stimulation was retinotopically aligned (V1 median AOK = −0.43, p = 0.04). This result suggests that inhibiting parts of V1 that are outside the stimulus representation may weakly facilitate detection performance, perhaps by quenching noise arising from task-irrelevant neurons. There was no modulation in the stimulus-offset SC AOK (median SC AOK = 0.01, p = 0.5). Moreover, the AOK distributions for offset sessions for V1 and SC were significantly different from one another (p < 0.0001; Kruskal-Wallis test). These data argue that the effects of white noise optogenetic stimulation, and thus the estimated NBK, is specific to the neurons in SC or V1 that represent the stimulus the animal is trying to detect. However, inhibiting nearby irrelevant areas of V1 may slightly facilitate performance compared to the equivalent manipulation of SC.

**Figure S4.**
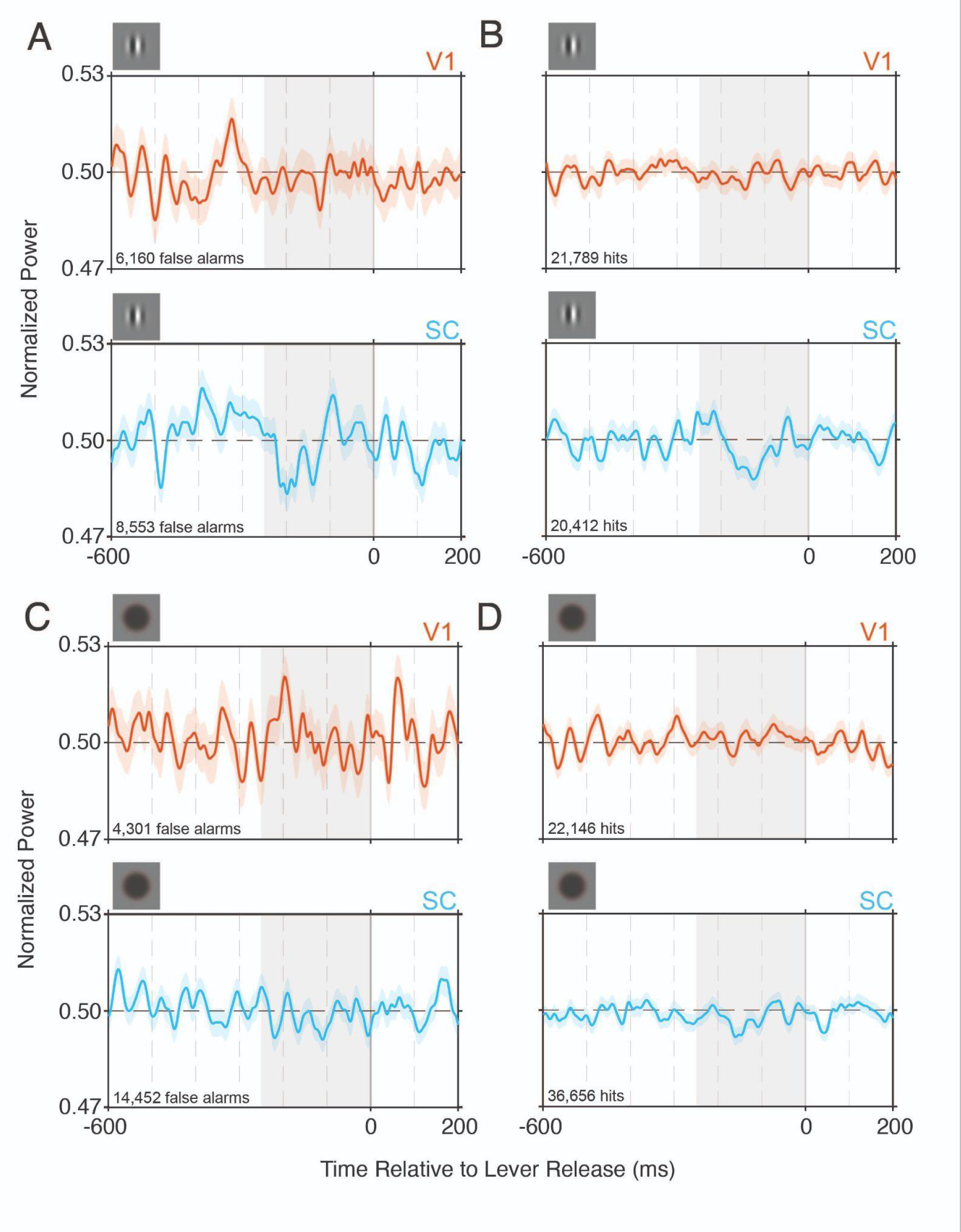
NBKs aligned to behavioral responses separated by visual stimulus conditions. These data are related to Figure 3 where the behavioral response aligned NBKs were combined across luminance and contrast sessions. Lever responses occur at t = 0. The analysis window used to compute the area over the NBK (AOK) is indicated by the gray box. Trial counts are indicated in the bottom left corner of each plot. Shaded area depicts ±1 SEM. **A)** (top) False alarm aligned NBK from V1 computed from contrast detection sessions (8 mice, 209 sessions). We computed the AOK from these traces during the 250 ms preceding the lever response using a bootstrap and found the distribution of AOK values was not significantly different from zero (median AOK = 0.69, p > 0.05). Consistent with the results shown in Figure 3, this argues that optogenetic stimulation of V1 did not promote false alarms. (bottom). False alarm aligned NBK from SC computed from contrast detection sessions (7 mice, 185 sessions). The false alarm aligned AOK for SC trended toward a difference from zero (median AOK = 1.36, p = 0.06). Thus, reductions in the optogenetic stimulus power may have promoted false alarms if the drop in power occurred ∼200 ms earlier. The timing of these effects is more consistent with inducing a phosphene or another fictive percept rather than directly promoting motor movements. **B)** (top) NBK from V1 aligned to lever releases on hit trials (where a visual stimulus was presented before t = 0) computed from contrast detection sessions (8 mice, 209 sessions). Consistent with V1s role in early sensory processing, there were no obvious modulations in the V1 NBK before lever releases and this was supported by the bootstrapped AOKs which were not significantly different from zero (median = 0.48, p > 0.05). (bottom) NBK from SC aligned to lever responses on hit trials for visual contrast (7 mice, 185 sessions). The AOK during the analysis window was significantly different from zero (median = 1.09, p = 0.01). The modulation in the NBK is broader than the stimulus aligned NBK from SC in Figure 2A and there is no sign of a modulation immediately before lever releases suggesting the primary role of SC in this task is related to sensory processing. **C)** (top) Same as in A except for luminance detection sessions with V1 stimulation (6 mice, 163 sessions). The AOK distribution from the V1 false alarm NBK was not significantly different from zero (median = −0.65, p > 0.05). (bottom) Same as in A except for luminance detection sessions with SC stimulation (11 mice, 367 sessions). Comparable to the data for contrast detection sessions, the AOK computed from SC false alarms was NBK was significantly different from zero (median = 1.08; p = 0.04). As above, this supports the idea that momentary reductions in the optogenetic stimulus power could produce a signal that the mice use to guide their responses. **D)** (top) Same as in B, except for V1 optogenetic stimuli aligned to behavioral responses on hit trials for luminance changes (6 mice, 163 sessions). The AOK distribution from the V1 hit-aligned NBK was not significantly different from zero (median = −0.74, p > 0.05). (bottom) Same as in B except for luminance detection sessions with SC stimulation (11 mice, 367 sessions). The AOK during the analysis window was significantly different from zero (median = 0.92, p < 0.05). The modulation in the NBK is broader than the stimulus aligned NBK from SC in Figure 2C and there is no sign of a modulation immediately before lever releases suggesting the primary role of SC in this task is related to sensory processing.

**Figure S5.**
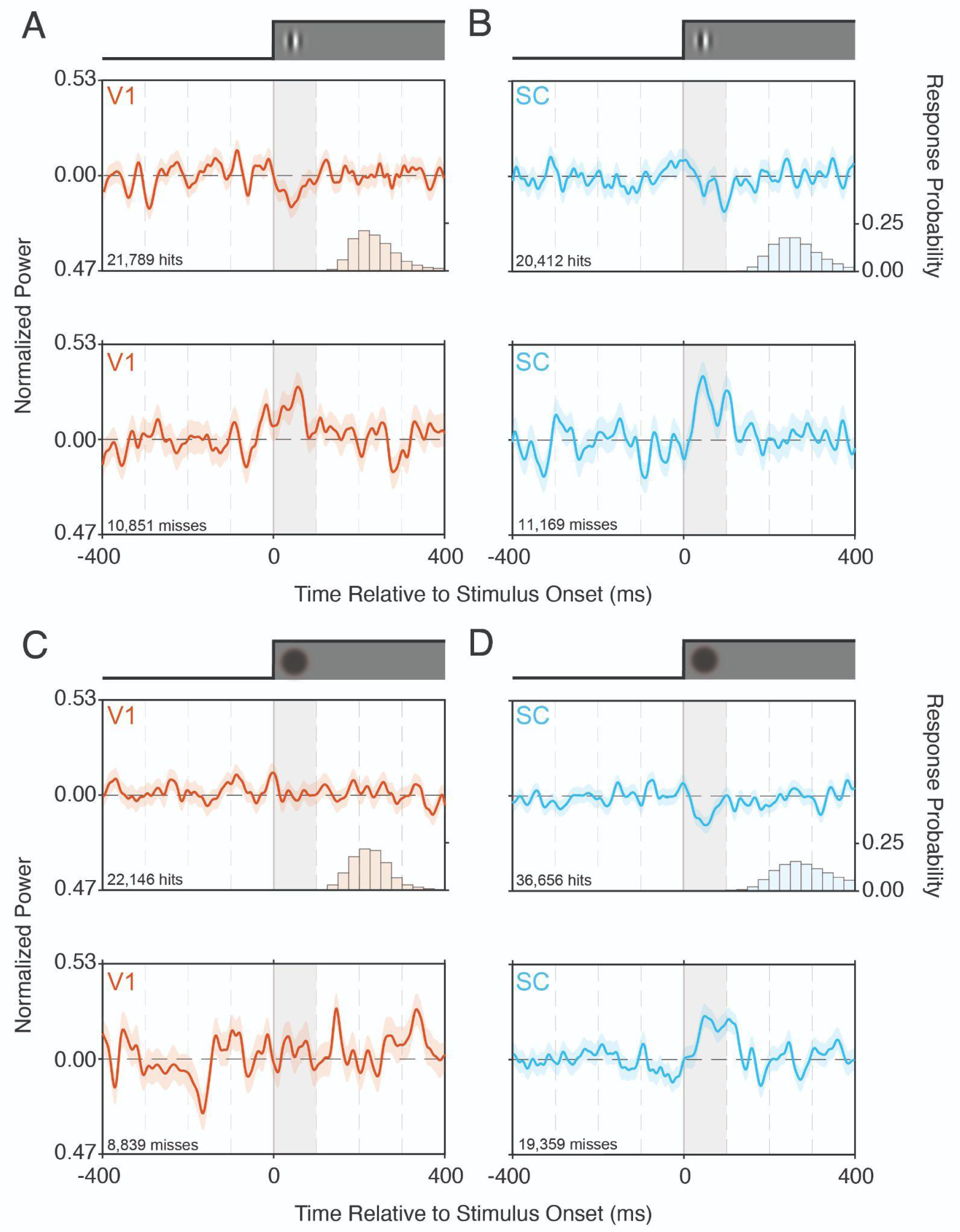
Hit and miss kernels separated by brain area and stimulus type. Data are related to Figures 2-3. Visual stimulus onset occurs at t = 0. The analysis window used to compute the area over the NBK (AOK) shown in Figure 2 is indicated by the gray box. Trial counts are indicated in the bottom left corner of each plot. Shaded area depicts ±1 SEM. The profile and type of visual stimulus is depicted above each panel. **A)** (top) stimulus aligned NBK from V1 computed from hit trials during contrast detection sessions (8 mice, 209 sessions). The small negative deflection in the NBK during the first 100 ms after stimulus onset (t = 0) shows that successful detection of contrast changes are more likely when V1 optogenetic stimulation is reduced during this period. (bottom) stimulus aligned NBK from V1 computed from miss trials during contrast detection sessions. The positive deflection during the initial 100 ms after the stimulus onset shows that misses are more likely to occur when the optogenetic power is elevated during this period. **B)** Same as in A) except for sessions with SC stimulation (7 mice, 185 sessions). **C)** Same as in A) except for sessions in which white noise optogenetic inhibition was delivered to V1 during luminance detection sessions (6 mice, 163 sessions). There were no obvious modulations in either the hit (top) or miss (bottom) NBKs around the time of stimulus onset. **D)** (top) stimulus aligned NBK from SC computed from hit trials during luminance detection sessions (11 mice, 367 sessions). The negative deflection in the NBK during the first 100 ms after stimulus onset (t = 0) shows that successful detection of luminance changes are more likely when SC optogenetic stimulation is reduced during this period. (bottom) stimulus aligned NBK from SC computed from miss trials during luminance detection sessions. The positive deflection during the initial 100 ms after the stimulus onset shows that misses are more likely to occur when the optogenetic power is elevated during this period.

**Figure S6.**
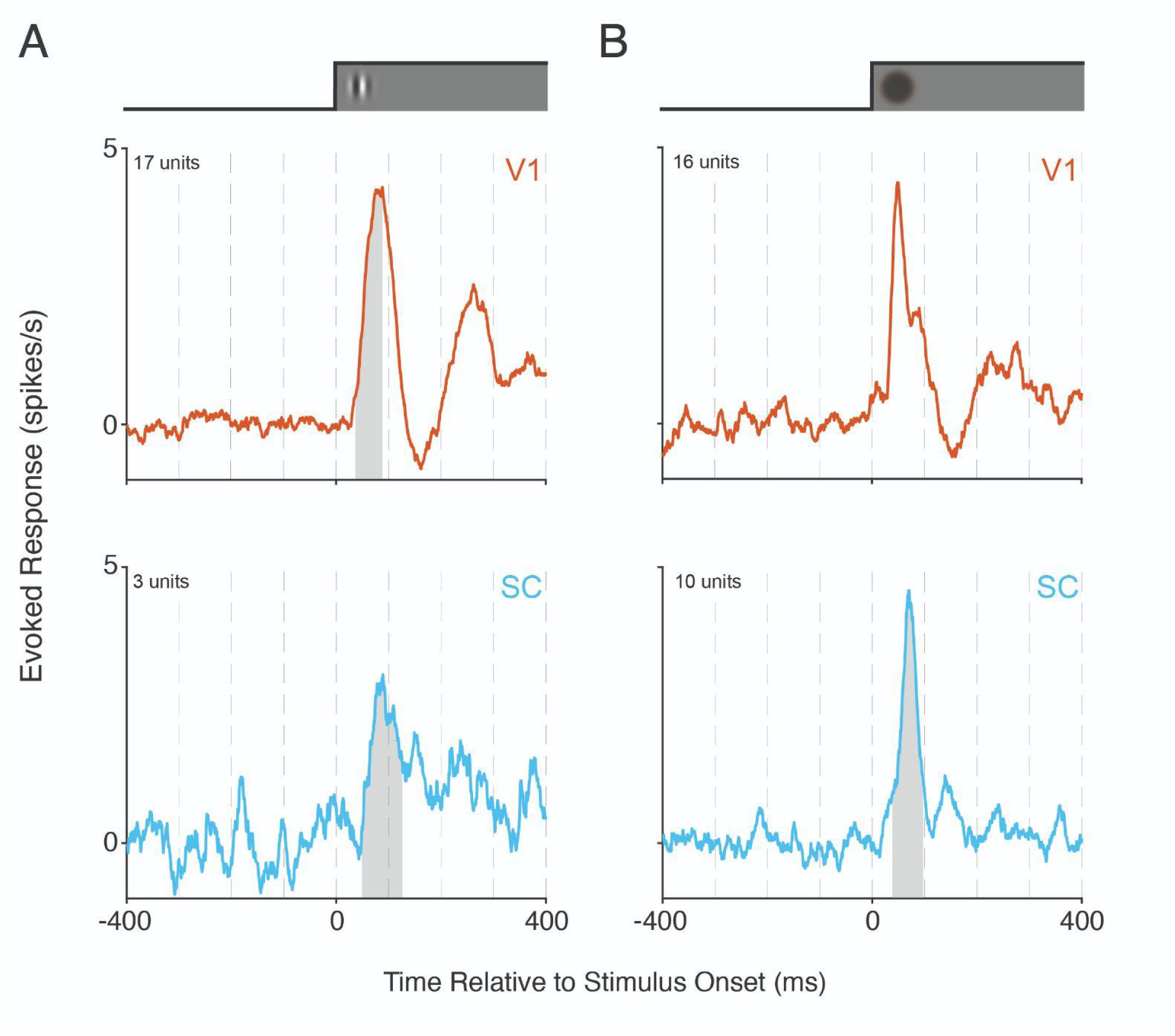
V1 and SC units respond to contrast and luminance changes. We recorded from many neurons simultaneously, such that visual stimuli were suboptimal for the majority of recorded units. To assess visual responsiveness, we compared the firing rate of each neuron during the 10-110 ms following the onset of a visual stimulus with the equivalent pre-stimulus period only during trials in which optogenetic stimulation was omitted. The analysis window was chosen to match the window applied to the NBKs in Figure 2, but shifted by the 10 ms delay to changes in spiking discovered from the STA analysis in Figure 4. A small proportion of neurons in V1 and SC exhibited significant visual responses to contrast or luminance changes (V1 contrast: 20/93 units; SC contrast: 3/78 units; V1 luminance: 19/113 units; SC luminance : 12/112 units; p < 0.05, Wilcoxon signed-rank test). The vast majority of visually responsive units were excited by changes in visual input regardless of brain area or stimulus type. The plotted above traces correspond to the average evoked response (spikes/s - baseline spike rate) across the significantly excited units (V1 contrast: 17/20, top left, red trace; SC contrast: 3/3, bottom left, blue trace; V1 luminance: 16/19, top right, red trace; SC luminance: 10/12, bottom right, blue trace). The profile of the visual stimulus is depicted directly above the plot. The gray shaded region depicts the FWHM of the corresponding NBK shifted by the 10 ms delay to inhibition determined from the STAs in Figure 4. The gray region is omitted from the V1 luminance response (top right) because there were no obvious modulations evident in the V1 luminance NBK. In sum, we encountered units that responded to both stimulus types in both SC and V1 and the majority of these units were excited by visual input.

**Figure S7.**
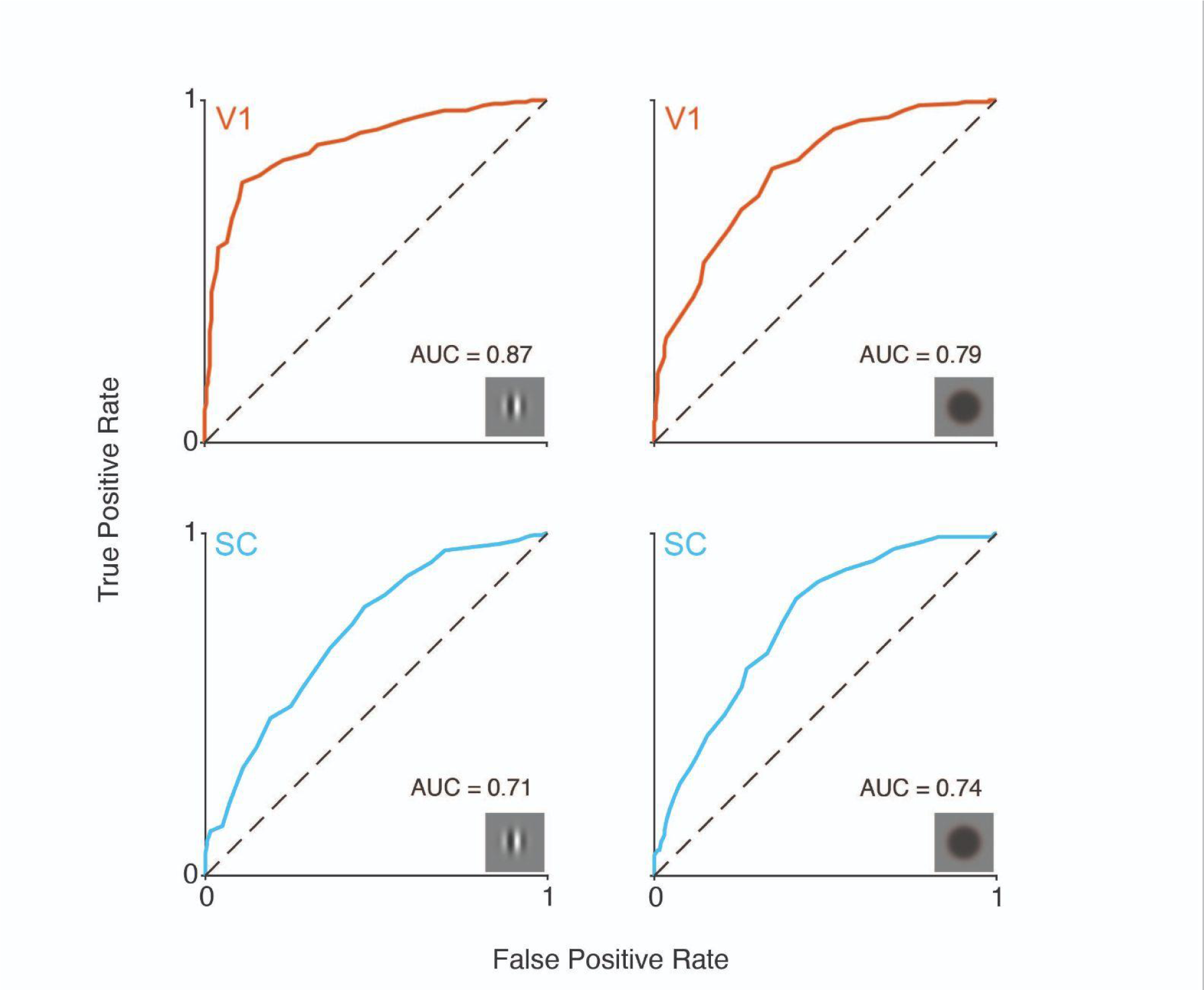
ROC analysis for population spike counts before vs. after stimulus onset. ROC curves for V1 (red) and SC (blue) spike counts during the 10-110 ms after visual stimulus onset compared to during the equivalent pre-stimulus window. This analysis only considered the sub-populations of neurons for each brain area or stimulus type that were significantly excited by visual stimuli (i.e., the same neuronal populations plotted in Figure S6). The gray box in the lower right corners indicates the stimulus type. Notably, the auROC values indicated that the onset of visual stimuli could be reliably discriminated from baseline spike counts regardless of brain area or stimulus type (auROC: V1 contrast: 0.87; SC contrast: 0.71; V1 luminance: 0.79; SC luminance: 0.74). These data suggest that even though there is information that could be used to detect luminance changes in V1, the lack of a strong modulation in the V1 NBK for luminance argues that mice don’t take advantage of these signals for guiding their behavioral responses.

**Figure S8.**
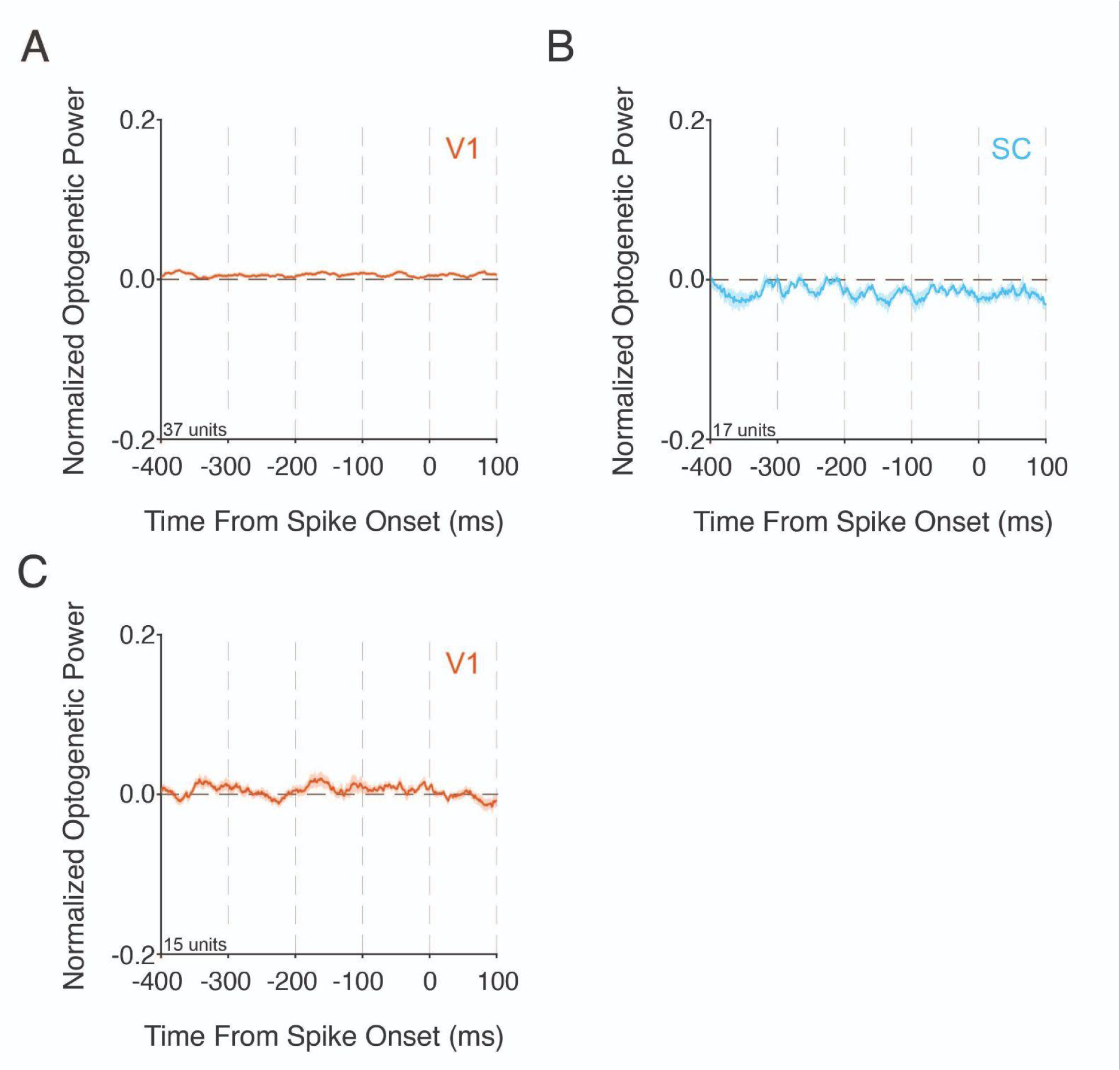
STAs from control recordings. SC and V1 are reciprocally connected via direct and indirect connections,^52–54^ and changes in spiking in V1 (SC) can impact processing in SC (V1).^55,56^ We thus wanted to determine whether white noise optogenetic perturbations of SC disrupted processing in V1 (and vice versa). We performed electrophysiological recordings in V1 or SC while we delivered white noise optogenetic stimulation to the other brain area to compute cross-area STAs. Prior to recording sessions, all mice were trained to respond to visual stimuli (see Materials and Methods) and then injected with AAVs. Mice then underwent initial optogenetic stimulation sessions to optimize retinotopic alignment between visual and optogenetic stimulation (see Materials and Methods). During recording sessions, the receptive field location of the recording electrode was optimized to overlap with the site of maximal optogenetic effect in the other structure identified in the behavioral alignment sessions. **A)** We recorded 37 units from 10 V1 unique recording locations in 3 mice (all male) while delivering white noise optogenetic stimulation to ChR2 expressing SC neurons. We then aligned the SC optogenetic stimuli to V1 spike times to compute a V1 population STA conditioned on optogenetic input to the SC. The V1 STA conditioned on SC white noise optogenetic stimuli shows no obvious modulations in the optogenetic power at any time prior to V1 spikes, suggesting that the white noise optogenetic stimulation in SC did not appreciably augment V1 spiking. **B)** Same as in A, except for V1 white noise optogenetic stimuli aligned to SC spike times. We recorded 17 SC units across 5 unique SC locations in 2 mice (1 female) while delivering white noise optogenetic stimulation to ChR2 expressing V1 neurons. Similar to above, the SC population STA conditioned on V1 white noise optogenetic stimuli revealed no obvious modulations in the V1 optogenetic stimulation power prior to SC spikes. Together, these data support the idea that while V1 and SC share reciprocal direct and indirect connectivity, white noise optogenetic stimulation delivered to V1 or SC at the powers used during behavioral sessions did not disrupt spiking in the other structure, suggesting the effects of our perturbations on performance depended primarily on the structure receiving optogenetic stimulation rather than via indirect effects. **C)** Given that white noise optogenetic stimulation involves repeated stimulation both within and across trials, we asked whether the effects we observed resulted from non-specific disruption of neuronal activity caused by prolonged energy transfer to the brain (e.g., heat). We conducted additional electrophysiological recordings at 4 locations in V1 of 2 mice (1 female) in which we delivered white noise optogenetic stimulation concurrently with visual input (see Materials and Methods). These mice were not previously injected with ChR2 AAVs. Furthermore, the optogenetic power was doubled compared to all other STA recordings (mean power = 0.5 mW) to ensure that energy transfer to the brain would exceed that seen in behavioral experiments. We recorded responses from 15 units in V1 and computed the population STA. There were no obvious modulations in the V1 population STA at any time proceeding spikes. These data suggest that modulations in the STA (and NBK) result from direct activation of opsin expressing neurons and not from indirect changes in activity due to tissue heating. Shaded area depicts ±1 SEM. Note the y-axis range is equivalent to those in Figure 4B to ensure direct comparisons across STAs.

